# It’s Not Always Crystal Clear: In-Pocket Analysis for Understanding Binding Site Microenvironment Properties

**DOI:** 10.1101/2025.09.24.678379

**Authors:** Filipe Menezes, Tony Fröhlich, Lucy C. Schofield, Julian Cremer, Jon Agirre, Robbie P. Joosten, Raphael Bourgeas, Sihyun Sung, José Antonio Marquez, Garib Murshudov, Oliver Plettenburg, Michael Sattler, Grzegorz M. Popowicz

## Abstract

Structural biology enabled key advances in biochemistry, drug discovery, and AI-driven three-dimensional structure modelling. Despite its importance, reliable and general computational tools for chemical validation of structural models remain limited, especially when small-molecule ligands are involved. This challenge is exacerbated when experimental resolution is insufficient to define interaction chemistry, leaving critical interpretations ambiguous. We introduce In-Pocket, a quantum mechanical optimisation method biased by experimental data that evaluates the chemical nature of protein-ligand binding interfaces. In-Pocket improves model quality, corrects chemically inconsistent structures, and identifies the energetically most favourable ligand-conformers within biomacromolecular complexes. The approach is generally applicable to a wide range of systems, including those containing transition metals. Benchmarking demonstrates that In-Pocket enhances the reliability of legacy and newly solved structures. With automated, user-friendly routine chemical validation, In-Pocket offers unique opportunities to improve the integrity of structural databases and impact diverse domains: from molecular biology to rational drug design and application of generative AI.

## INTRODUCTION

The Protein Data Bank (PDB)^1^ is the reference repository for biological macromolecular structures. This foundational community resource comprises over 235,000 entries and has enabled the development of transformative machine-learning (ML) approaches for protein structure prediction.^2,3^ The majority of PDB structures have been determined by macromolecular X-ray crystallography (MX),^4^ with additional contributions from nuclear magnetic resonance (NMR), and a rapidly growing number now determined by single-particle cryo-electron microscopy (cryo-EM).^5^

All current experimental structure-determination methods are fundamentally constrained by resolution. Outside rare sub-Ångström cases, typical MX and cryo-EM datasets exhibit limited precision in resolving bond lengths and angles within heteroatom scaffolds,^6,7^ lack direct experimental evidence for hydrogen atoms, and often present ambiguities in density interpretation. In addition, commonly used representations of molecular scaffolds rely on simplified valence models that do not explicitly encode electronic structure. Inaccurate ligand structural assignments, in which bond definitions dictate the geometry and strength of intermolecular interactions, can therefore propagate downstream errors in chemical interpretation. These limitations affect the reliability of structural models, particularly in applications that require accurate binding energetics, such as structure-based drug discovery and the development of data-driven ML models. As a result, chemically informed computational validation of biomolecular structures remains an important and incompletely addressed challenge in structural biology.

To address this challenge, we introduce In-Pocket Analysis (IPA), an algorithm for evaluating the local chemical environment of a compound or residue within a macromolecular structure (**Fig. 1a-d**). Although frequently applied to small-molecule ligands, IPA is equally applicable to amino acids and their modifications, nucleic acids, and metal-containing cofactors. The method performs experimentally biased quantum-mechanical (QM) optimisation of ligand-environment interactions across multiple protonation and tautomeric states, referred to here as chemical states. By validating against deposited experimental structures, we demonstrate that IPA enables a quantitative assessment of ligand chemical states and geometries within their structural context.

**Figure 1:**
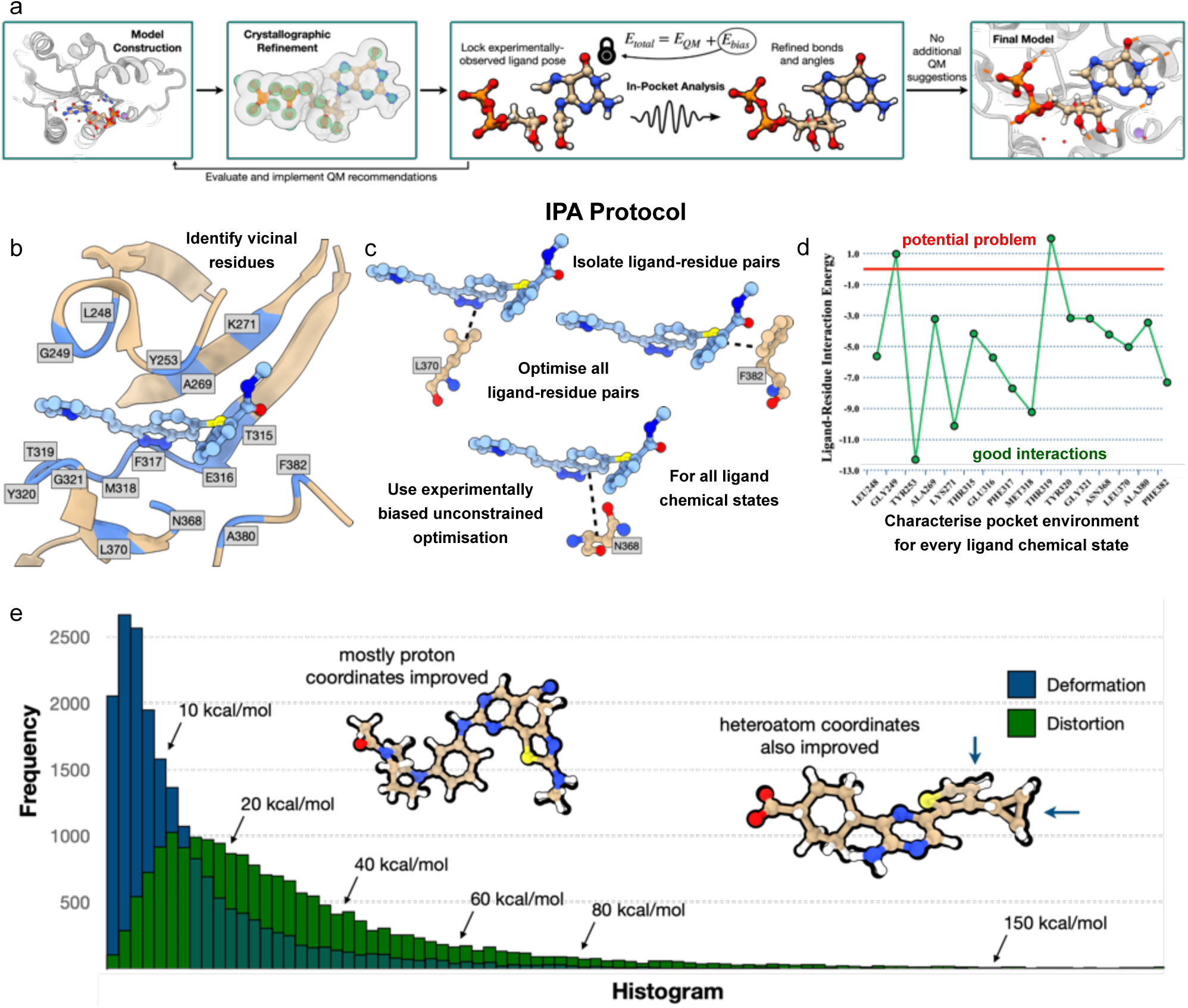
IPA for detailed chemical examination of biological macrocomplexes. a-d) The IPA protocol. a) Modified crystallographic model-solving pipeline with IPA embedded for chemically refined structural models. Given a biological structural model, b) IPA identifies all residues within the vicinity of the active residue, typically a small molecule ligand. c) IPA then uses a series of experimentally biased QM optimisations of each molecular pair to d) calculate experimentally relevant interaction energies. When negative, these energies indicate good, attractive interaction pairs. On the other hand, positive interaction energies indicate possible pathologies in the structural model. e) IPA is used to analyse experimental model distortion and the ligand deformation energy of 20,000 ligands available in binding databases.^8,9^ Two histograms are obtained: a distribution peaked at about 5.0 kcal.mol^−1^ with a rapid decay is observed for IPA-derived ligand deformation energies. A broader distribution peaking at 10-20 kcal.mol^−1^, slower decay and an extremely long tail is obtained for the experimental distortion energies.

## RESULTS

To analyse ligand-bound poses in protein-ligand complexes, IPA quantifies ligand-environment interaction energies (E_int_), ligand deformation energies (E_def_), ligand distortion energies (E_dist_), and relative energies of ligand chemical states (E_cs_). E_def_ represents the receptor-induced energetic cost required for a free ligand in solution to adopt its bound conformation, whereas E_dist_ reflects deviations from chemically expected ligand geometries in experimentally determined structures. These quantities are automatically evaluated across multiple ligand chemical states and combined into a binding energy term (E_bind_=E_int_+E_def_+E_cs_). This enables systematic comparison of ligand chemical states and geometries across structural models. Supplement **S1** provides a theoretical rationale for E_dist_ and its use in downstream analyses.

### Ligand Structural Distortion in Protein-Ligand Complex Databases

The binding of a ligand to a protein entails dovetailing interactions that reshape both partners and are associated with finite deformation energies. To quantify ligand distortion in experimentally determined protein-ligand structures, we used IPA to evaluate statistical metrics of distortion and deformation across a protein-ligand structure database (**Fig. 1e**).^8,9^ Ligand deformation energies exhibit a narrow distribution, peaking around 5 kcal.mol^−1^ and decaying rapidly. In contrast, the ligand distortion energy distribution is broader, with a maximum between 10-20 kcal.mol^−1^ and a long tail extending beyond 1,000 kcal.mol^−1^. Distortion energies are typically 2-4-fold larger than the deformation energies associated with ligand binding. Such deviations exceed physically expected deformation energies and are consistent with distortions arising from limitations in experimental resolution and modelling procedures.

The origins of ligand structural distortion are multifold. At low distortion energies, inaccuracies are primarily associated with proton placement and minor angular deviations. At intermediate energies, we observe short hydrogen-hydrogen contacts, pronounced angular distortions, and moderate bond-length deviations. For example, in compounds containing nitro groups, N–O bond lengths deviate by 0.1-0.2 Å from the reference value of 1.22 Å.^10,11^ At high distortion energies, ligands display severe bond-length and angular distortions, including simultaneously over-compressed and over-extended bonds within the same molecule. Such distortions are associated with misassignment of atomic hybridisation and protonation states when PDB-derived ligand topologies are neglected.^9^ Representative examples are discussed below (**Fig. 5**).

### Resolution of ligand protonation and tautomeric states in protein binding pockets

IPA’s most direct application is to determine a ligand’s optimal chemical state within a defined protein-ligand binding interface. After specifying a residue of interest–ligand, amino acid, or cofactor, IPA identifies all neighbouring residues within a defined distance threshold–*e.g.*, 4 Å–and it performs an experimentally-guided QM refinement to evaluate ligand-residue binding energetics (**Fig. 2a**). As an initial example, we analyse the crystal structure of axitinib bound to unmutated ABL1 (PDB-ID 4wa9; CCD-ID AXI, the ligand identifier).^4^ IPA identifies six chemical states, CS1-CS6, which were all considered in the analysis (**Fig. 2b**). Favourable binding geometries are characterised by the absence of strongly repulsive ligand-residue interactions, as those interactions are not physically motivated (**Fig. 2c**). Notably, CS1–which corresponds to the previously assigned but, based on this work, revised ligand topology–displays a proton-proton clash from the 2H-indazole tautomer (**Fig. 2d**). Deprotonating the indazole, as in CS3, destabilises interactions with the hinge region (F317-G321) due to unfavourable electronic clashes, an effect captured by the QM treatment employed by IPA. CS2 and CS4 correspond to high-energy tautomers of CS1 and CS3 and are energetically unfavourable; CS5 and CS6 exhibit favourable interaction landscapes among the six identified chemical states, since the 1H-indazole tautomer avoids steric and electronic clashes while forming two hydrogen bonds (**Fig. 2e**).

**Figure 2:**
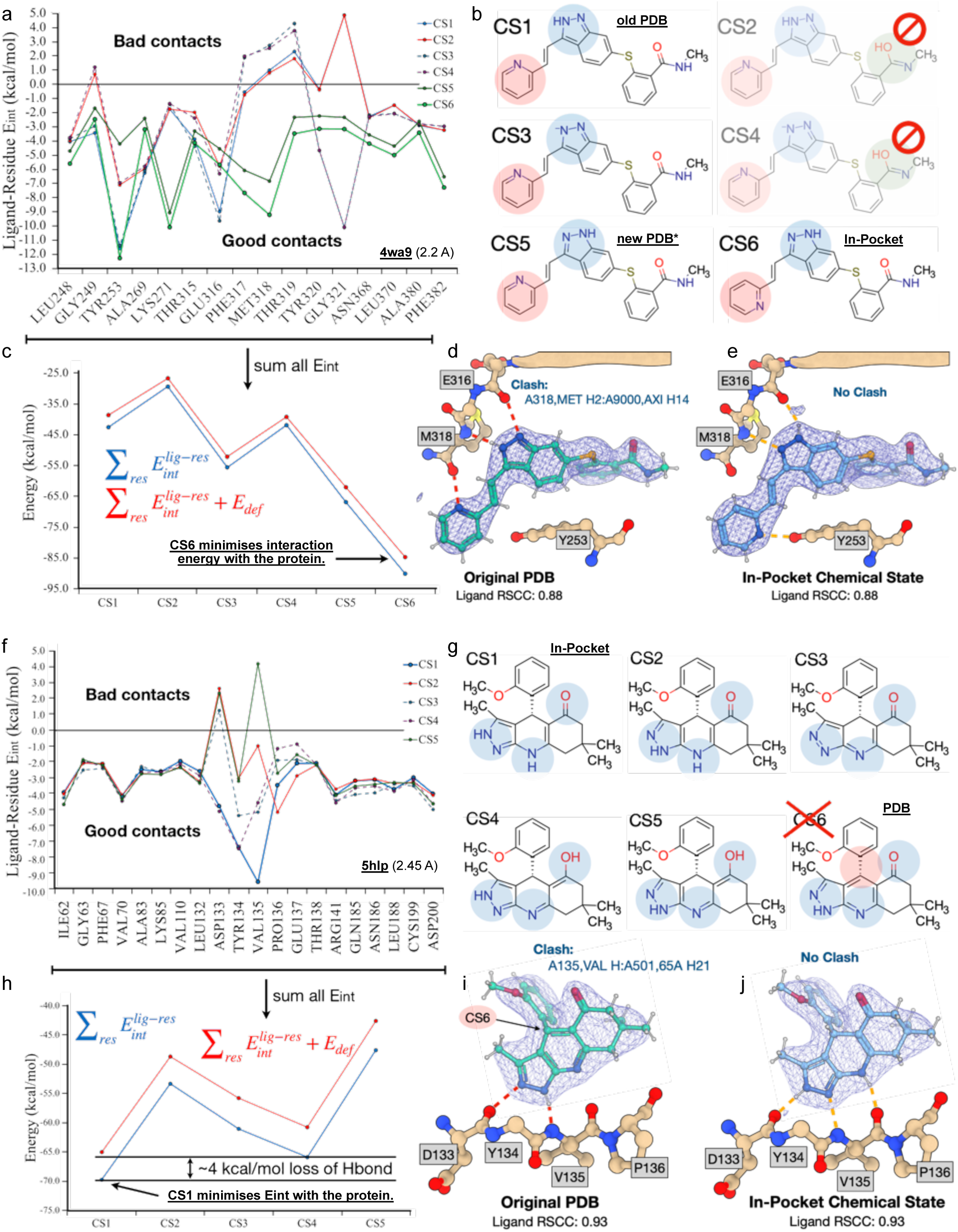
Investigating acid-base properties of ligands in a protein’s pocket: correcting ligand chemical states in kinase ATP-pockets. a-e) The binding analysis of axitinib in ABL1. f-j) Pyrazole equilibria in GSK3β. a) The ligand-residue interaction energy plot for axitinib in ABL1 for six axitinib chemical states identified by IPA. The calculations show that only chemical states CS5 and CS6 avoid suboptimal protein contacts. b) Axitinib chemical states identified by IPA. The chemical states CS2 and CS4 are faded because they are chemically unrealistic tautomeric states identified by the tautomer search; their energies are approximately 10 kcal.mol^−1^ higher than the respective amides (CS1 and CS3). The most favourable chemical state is CS6, which should be compared with the present PDB-deposited ligand topology (CS5) and the PDB-deposited chemical state before our reporting (CS1). The circles indicate which ligand functional groups differ between chemical states. c) IPA binding metrics to evaluate the best ligand chemical state in the protein’s pocket. The blue curve represents the summed ligand-residue interaction energies. This quantity is corrected in the IPA binding energy by adding the deformation energy (red curve). Irrespective of the metric, IPA selects CS6 as the most reasonable chemical state of axitinib in ABL1’s pocket. d,e) Experimental validation: IPA resolves the ligand-protein clash. d) The ligand’s tautomeric state before our reporting, and e) the new IPA-identified chemical state. The electron density at the experimental resolution is inconclusive. The depicted model representation also shows why CS6 is more favourable than CS5: by flipping the terminal pyridyl, this group forms an additional hydrogen bond with Y253. g) Ligand-residue interaction energy plot of a pyrazole-containing ligand in GSK3β’s pocket for the five IPA-identified ligand chemical states. CS2, CS3, and CS5 lead to poor protein-ligand contacts. h) IPA-identified ligand chemical states (CS1-5) and the PDB-deposited ligand chemical state (CS6). Note that the ligand topology declared in the PDB disagrees with what is reported in the original publication, indicating possible transfer issues. CS6 is disregarded in all calculations due to this inconsistency. i-j) Experimental validation using traditional crystallographic metrics. i) Schematic representation of the pocket and the main hinge residues interacting with the ligand and its density. The structural incompatibility between the PDB-deposited topology and the structural model is evidenced. j) Similar to block i) but for the IPA-selected ligand model. It can be followed that CS1 is more favourable than CS4 due to a hydrogen bond, which yields CS1 an additional 4 kcal.mol^−1^ stabilisation.

At a resolution of 2.2 Å, empirical densities cannot reliably distinguish between carbon and nitrogen atoms (**Fig. 2d-e**). Consequently, density-based interpretations are insufficient to resolve all atomic identities in a structure. IPA provides quantitative support for evaluating alternative functional-group orientations. This is particularly relevant for axitinib’s pyridyl group, differentiating CS5 from CS6. Flipping the pyridyl moiety forms an additional, unrecognised hydrogen bond with Y253 (**Fig. 2e**), further stabilising the ligand within the binding pocket. The IPA-identified chemical state, CS6, is further supported by clash analysis (see experimental validation data in **S2**).

In the PDB-ID 5hlp (CCD-ID: 65A),^12^ an indazole-containing ligand is bound to the GSK3β kinase (**Fig. 2f-j**). The PDB-deposited aromatisation of the central ring is inconsistent with a chemically reasonable geometry (CS6 in **Fig. 2i**); therefore, the proposed indazole is not supported, and we considered the ligand with a pyrazole group instead. Note that the PDB-deposited ligand topology differs from what is declared in the publication.^12^ IPA identifies two ligand chemical states close to the starting PDB topology, with a 1H-pyrazole tautomer: CS2 and CS5. These chemical states clash with the kinase’s hinge residues, L132-T138 (**Fig. 2i-j**). Deprotonating the pyrazole leads to better interaction networks, as in CS3. However, the ligand’s pocket representation is further improved by considering the 2H-pyrazole tautomer, as in CS1 and CS4 (**Fig. 2j**). Among these states, CS1 provides the most favourable interaction profile in GSK3β, followed by CS4. Although the difference in binding energy is modest (4.0 kcal.mol^−1^), CS1 is favoured owing to an additional hydrogen bond compared to CS4. This assignment aligns with the publication,^12^ though a methyl group is still missing from the structure. The crystallographic refinement^13,14^ of 5hlp with the IPA-selected chemical state evidences the resolved clash with V135.

Further analyses of ligand-chemical states in protein pockets are provided in supplements **S2-S6**.

### Effects of ligand chemical state on protein-ligand binding thermodynamics

To assess the thermodynamic consequences of ligand chemical-state assignment, we examined axitinib binding to two ABL1 variants (**Fig. 3a**): the unmutated structure analysed above and ABL1^T315I^ (PDB ID 4twp)^4^. The IPA-selected axitinib chemical state is compared with the ligand topology originally deposited in the PDB. For reference, the IPA-selected chemical state is consistent with the crystal structure of ABL1^T315I^ and yields improved metrics in standard crystallographic validation protocols.

**Figure 3:**
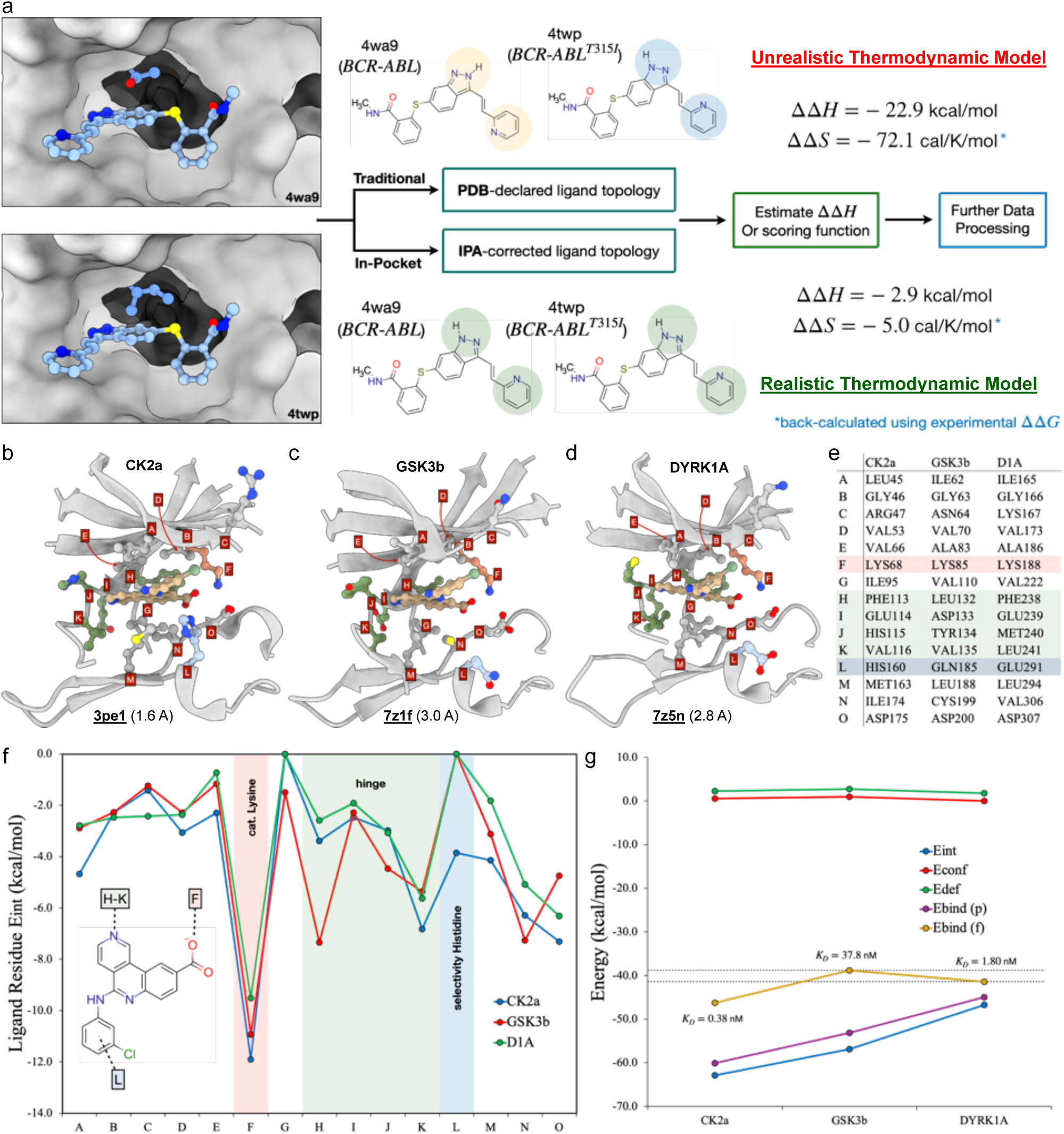
IPA for detailed chemical examination of biological macrocomplexes and translation to drug discovery applications. a) The impact of ligand topology is evaluated using two ABL1 variants in complex with axitinib: top left, unmutated protein–PDB-ID 4wa9; bottom left, the T315I mutant–PDB-ID 4twp. The relative QM binding energy should correlate with the relative enthalpy of binding from a hypothetical isothermal titration calorimetry (ITC) experiment. The relative Gibbs free energy is estimated from the IC_50_ ratios, and the relative binding entropies are back-calculated using ΔΔ” = (ΔΔ% − ΔΔ’)/298.15. Unrealistic binding thermodynamics data are obtained following the PDB-deposited ligand topology, whereas realistic thermodynamic parameters are obtained using the IPA-suggested ligand chemical state. b-g) Showcasing IPA’s guiding design capabilities in the complex problem of kinase selectivity. IPA is performed on protein-ligand complexes of silmitasertib (CCD 3NG) with b) CK2α (PDB-ID 3pe1),^16^ c) GSK-3β (PDB-ID 7z1f),^17^ and d) DYRK1A (PDB-ID 7z5n).^17^ To ensure comparability, residues in the pocket were positionally aligned, and only the conserved amino acids were considered for the analysis. e) The list of positional equivalencies in the amino acids of the three kinases. f) Ligand-residue interaction energies for the three kinases analysed. Three different regions were selected in the pockets, namely the catalytic lysine, the hinge regions, and a previously identified element of selectivity that grants silmitasertib its sub-nanomolar affinity to CK2α. g) Overall IPA metrics for the kinase selectivity example. The pairwise IPA binding energy, though useful for analysing model integrity, lacks cooperativity effects needed to score ligands in the pockets. This is recovered using IPA’s fragment mode, which includes a whole chemical environment.

We evaluated the impact of ligand chemical state on thermodynamic predictions using two modelling protocols: one that employs the PDB-deposited ligand topology and the other that uses the IPA-derived chemical state. QM energy calculations were used to estimate relative binding energies, which correlate with binding enthalpies. Models incorporating IPA-derived chemical states yielded relative energies closer to values commonly reported in isothermal titration calorimetry experiments, which directly measure binding enthalpy and entropy contributions. To further assess consistency with experimental observables, relative IC₅₀ values^4^ were used to estimate relative binding affinities, from which relative entropic contributions were inferred. These entropy estimates further strengthen the observation that IPA-ligand topologies yield thermodynamically consistent data.

### Energetic determinants of kinase selectivity

Kinase selectivity remains a central challenge in the development of protein kinase inhibitors.^15^ Achieving true selectivity requires a detailed understanding of drug-target interaction networks. To evaluate whether IPA captures energetic features underlying selectivity, we analysed the binding of silmitasertib, a sub-nanomolar CK2α inhibitor,^16^ across two additional kinase targets, GSK-3β and DYRK1A (**Fig. 3b-g**).^17^

**Fig. 3b-d** depict the structural models used for analysis (CK2α, PDB-ID 3pe1;^16^ GSK-3β, PDB-ID 7z1f;^17^ DYRK1A, PDB-ID 7z5n;^17^ CCD-ID 3NG). To directly compare the different binding sites, only positionally equivalent residues were considered (see residue mapping in **Fig. 3e**). Three regions were selected for detailed comparison (**Fig. 3f**): the catalytic lysine, the hinge segment, and the selectivity-histidine region.^17^ Among these, interactions involving the catalytic lysine are stronger in CK2α, followed by GSK-3β. In contrast, hydrogen bonding to the hinge region is stronger in GSK-3β, particularly at Leu132, suggesting a potential avenue for selective GSK-3β inhibitor design. CK2α shows the second strongest hinge contact at Val116, whereas hinge interactions are weaker in DYRK1A. The third region, corresponding to the selectivity residue *L*, shows a markedly stronger interaction with CK2α’s His160, consistent with its role as a key determinant of silmitasertib selectivity.^17^

Global binding metrics further highlight the importance of capturing local cooperative effects (**Fig. 3g**). IPA’s pairwise binding energy metric (E_bind_(p)) fails to reproduce the experimental affinity trend across these kinases. This reflects the absence of cooperative interactions in purely pairwise interaction descriptors. In contrast, reconstructing the same pockets in IPA’s fragment mode (E_bind_(f)) restores local environmental effects and correctly reproduces the experimental selectivity ranking.

### Resolving alternative structural models of protein-ligand complexes

Multiple structural models may exist for the same protein-ligand complex, either available across different PDB entries or as alternative conformations within a single model. These representations may conflict at the ligand-binding site or represent alternative binding poses in equilibrium. An illustrative example is curcumin bound to the DYRK2 kinase: in the PDB-ID 6hdr (CCD-ID CC9), curcumin adopts a binding mode where its central diketone group engages with the kinase’s hinge region (**Fig. 4a**), whereas in 5ztn (CCD-ID CUR),^18^ the diketone moiety is oriented away from the hinge (**Fig. 4b**). This discrepancy motivates evaluating whether one model provides a more accurate description of the biological complex or if both represent equally valid conformations.

**Figure 4:**
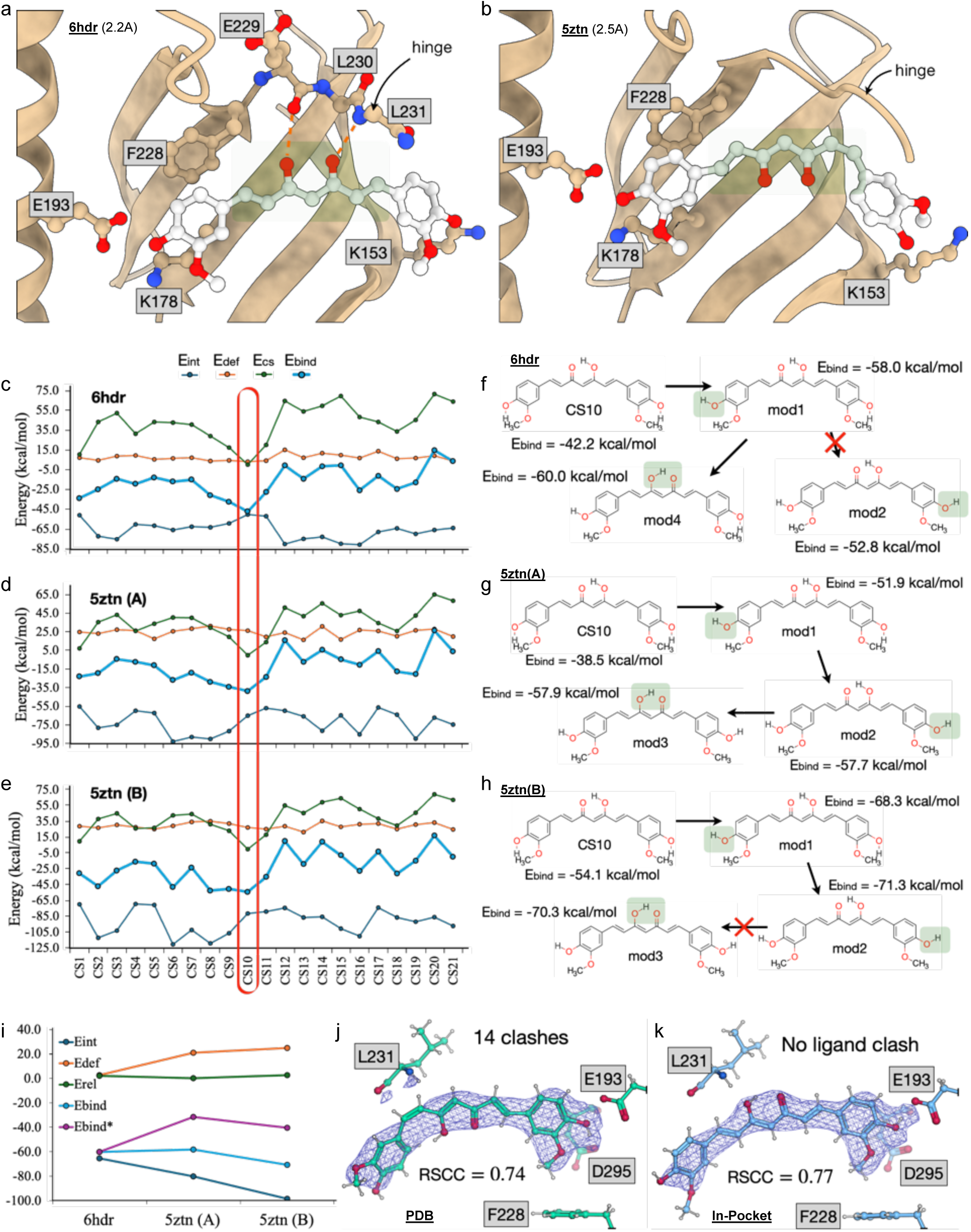
Understanding ambiguities in different protein-ligand structural models and using IPA for an informed selection of the best-representing pocket models: curcumin binding to DYRK2. a) The binding mode of curcumin to DYRK2 in the PDB-ID 6hdr. The two curcumin oxygens from the central diketone/keto-enol scaffold are within the interaction range of the hinge residues E229 and L231. b) The binding mode of curcumin to DYRK2 in the PDB-ID 5ztn, chain A. The central diketone/keto-enol group is flipped when compared to the 6hdr model, indicating that curcumin-hinge interactions are only lipophilic in nature, and no hydrogen bond to the hinge can be formed. There is additionally a change in ligand isomeric state, from trans-trans in 6hdr to trans-cis in 5ztn. c-e) The IPA binding metrics for curcumin in DYRK2. Besides the summed ligand-residue interaction energies and the deformation energy, the IPA binding energy also considers the relative energy of each chemical state to account for slight changes in the relative tautomer energies and the different protonation states. The list of all 21 IPA-identified ligand chemical states is provided in the supplementary material (**Figure S7**). The IPA binding metrics identify CS10 as the most favourable curcumin chemical state in DYRK2’s pocket, irrespective of the structural model. f-h) Guided hydrogen-atom network optimisation, as the initial guess proton coordinates might trap the system in metastable local minima: f) The refinement for the 6hdr model, where two changes in the initial proton coordinates are needed; g) The refinement for 5ztn’s chain A, where three changes in proton initial coordinates are required; h) The refinement of 5ztn’s chain B model, where two changes in the initial proton coordinates are needed. i) Overall IPA binding metrics to compare the structural models 6hdr and 5ztn. To compare the ligand’s binding pose in different pockets, a different reference metric is needed, where the change in the isomeric state must be considered. The final metric, Ebind*, represented in purple, clearly shows that 6hdr offers the best pocket representation. j) The 5ztn model validation, as deposited in the PDB. k) Reevaluation of 5ztn’s model validation using IPA’s selected chemical state and suggested binding mode. The model’s fitness has improved.

Applying IPA, 21 curcumin chemical states were identified (**Fig. S7**), all of which are compatible with the experimental data. Each state was evaluated within the three available representations: one chain from 6hdr and two chains (A and B) from 5ztn. Across all cases, IPA consistently identified CS10 as the most stable chemical state in DYRK2’s binding pocket (**Fig. 4c-e**). This suggests that curcumin’s protonation state is conserved across the differing ligand-binding poses. IPA binding energies incorporate interactions with the entire protein environment and chemical-state relative energies, accounting for factors such as tautomeric equilibria and changes in protonation states. Interestingly, CS10 disagrees with the ligand topology proposed in 6hdr–CCD-ID CC9–but aligns with that reported in 5ztn–CCD-ID CUR.

To further refine the ligand-residue interaction network, hydrogen-bonding patterns involving the phenolic groups and the central keto-enol moiety were systematically explored while maintaining the CS10 chemical state across all models (**Fig. 4f-h**; **Fig. S8**). Reorientation of the phenol group adjacent to E193 (**Fig. 4f-h**, mod1) reduced the binding energy across all three models (**Figure S8**). Rotating the phenol group near K153 (**Fig. 4f-h**, mod2) stabilised the 5ztn models but was unsuccessful for 6hdr. Finally, recognising that a proton is stabilised between the two oxygens of curcumin’s keto-enol group, proton transfers between these oxygens were explored (**Fig. 4f-h**, mod3 and mod4; **Fig. S8**), resulting in lower binding energies in 6hdr and 5ztn(A) but not in 5ztn(B).

While these refinements provide a chemically accurate description of curcumin within each structural model, direct inter-model comparisons remain unfeasible because the 6hdr structure features a trans-trans curcumin isomer, whereas 5ztn contains a cis-trans isomer. Further, residual experimental noise varies across models even after IPA refinement. To enable direct comparison, a renormalisation term was introduced to balance the relative energies of refined models (Ebind* in **Fig. 4i**). This correction disfavours the 5ztn models, ultimately supporting 6hdr as the most chemically plausible representation of curcumin binding to DYRK2. Consistent with this interpretation, refinement of the 5ztn model according to IPA-derived suggestions (**Fig. 4j-k**) removes ligand-associated clashes and improves the real-space correlation coefficient (RSCC),^19,20^ as assessed by standard crystallographic validation metrics.

Additional examples addressing common crystallographic ambiguities, including transition-metal and crystal-contact effects, are provided in the supplementary material. This includes analyses of hydrolysed imipenem bound to a zinc-containing β-lactamase (PDB-ID 6rzr,^21^ **S9**) and of a venetoclax-BCL-2 complex (PDB-ID 6o0k,^22^ **S10**).

### Correction of experimentally induced distortion in protein-ligand structures

Structure-determination pipelines can introduce significant distortion into structural models, impacting chemical interpretation. Representative examples are shown in **Fig. 5**. For example, an MDM2 inhibitor (PDB-ID 4mdn^23^, CCD-ID Y30) exhibits a distorted formamide group with ambiguous topology (**Fig. 5a**). To assess whether IPA improves model quality, we refined the structure using the PDB-deposited ligand topology (**Fig. 5b**). In the initial model, the ligand displayed an unrealistically high deformation energy of 1,710.6 kcal.mol^−1^, which dropped sharply to 79.7 kcal.mol^−1^ after one IPA-refinement iteration (**Fig. 5c**). This first step accounts for the largest distortion correction.

**Figure 5:**
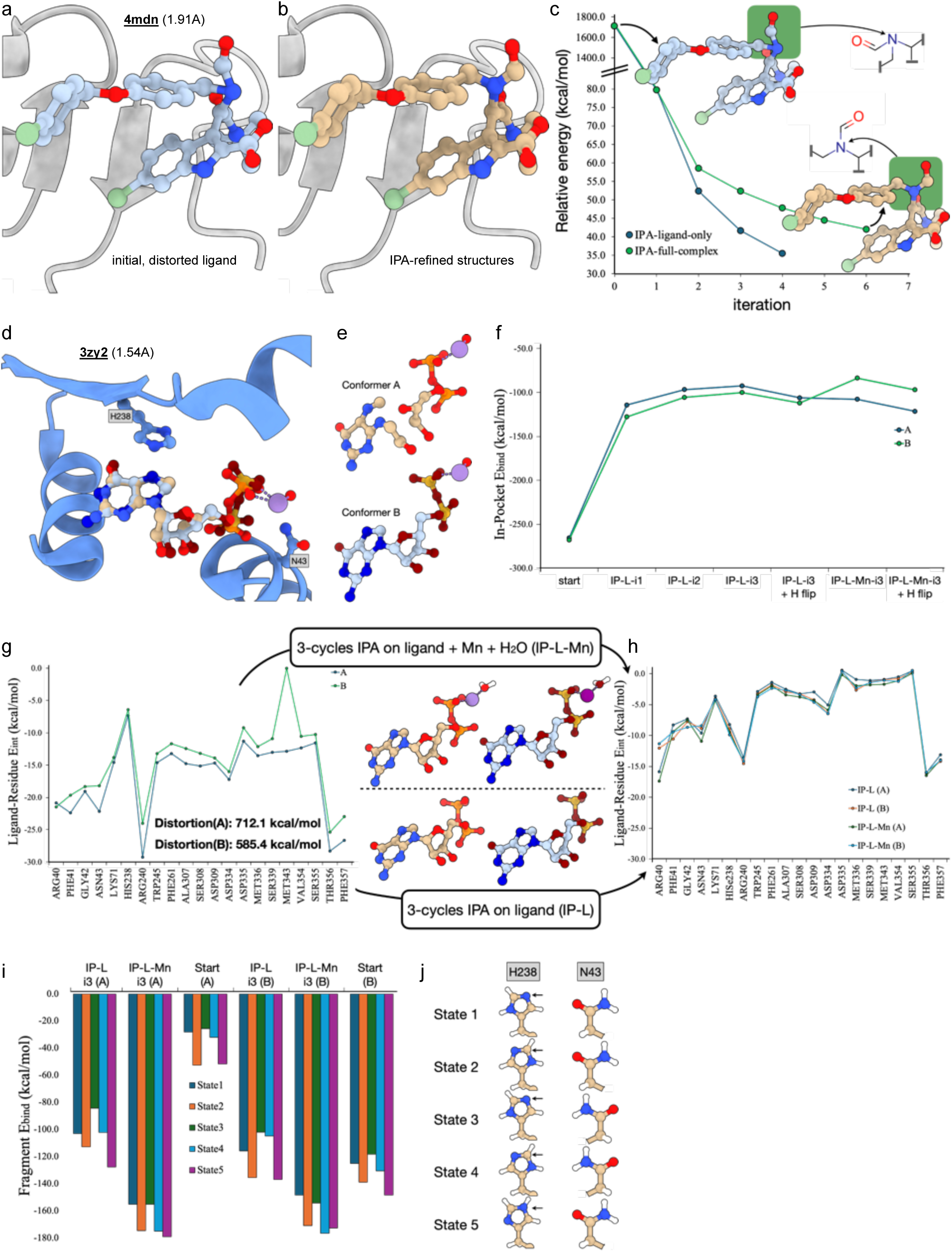
IPA mitigates structural distortions. a) PDB-deposited structure of 4mdn. The formamide group adopts a distorted conformation that deviates from the theoretically expected 120° angle, due to sp^2^ hybridisation. b) The IPA-fixed structure of the ligand Y30 shows the formamide group’s improved conformation. Two structures are superimposed: one in which IPA was applied to the isolated ligand, the other to the whole protein-ligand complex. The structural differences between ligands are minimal. c) Ligand relative energy as a function of IPA refinement iterations. Two refinement strategies were followed after the first ligand-only iteration: we optimised the ligand in isolation (blue) and the ligand within the protein’s pocket (green). Though the ligand’s geometry is fixed in both cases, we observe a slight drift in the ligand-only optimisation. d) The 3zy2 PDB structure shows two alternate conformations of guanosine diphosphate (GDP) bound to the POFUT1 fucosyltransferase 1. e) The three-dimensional representation of the two alternate conformations of GDP in POFUT1’s pocket, showing severe structural distortion. There are bonds missing because the representation software failed to recognise covalent linkages due to the severe structural distortion. f) IPA binding metrics for the two alternate ligand conformations. The ligand’s overstabilisation in the pocket observed for the PDB structures results from accumulated distortion. This has already been significantly improved with one IPA cycle. The initial PDB structure and the ligand-only IPA-refined structures indicate the degeneracy of the two alternate conformations. The degeneracy is lifted when the Mn^2+^ and the water molecule are included in the IPA ligand refinement cycles. On the abscissa, “start” indicates the initial structure extracted from the PDB; “IP-IL-i1”-“IP-IL-i3” indicates the ligand-only IPA refinement cycles (cycle number indicated last); “IP-IL-i3 + H flip” is identical to “IP-IL-i3” but with flipped H238; the last two points are the result of three IPA refinement cycles where the Mn^2+^ ion was included as part of the ligand, without and with H238 flip. g) IPA’s ligand-residue interaction energy plot is severely influenced by the structural distortion of each alternate conformation of GDP in the pocket. The calculated distortion energies are considerable. h) IPA’s ligand-residue interaction energy plot after three cycles of IPA ligand refinement using only GDP (path below) and using GDP with a coordinated Mn^2+^ ion and a water molecule bound to the metal (path up). The resulting interaction energy plots are more sensitive and faithful in describing the ligand in the pocket. However, the results are not entirely equivalent. i) The binding energy for fragments extracted from the 3zy2 complex at different residue-orientation states, summarised in j). The lowest binding energies are obtained when the ligand is IPA-refined together with the Mn^2+^ ion and the bound water molecule. The results show that the alternate configurations bind preferentially to different protein conformation states.

Following this correction, two refinement strategies were explored: optimisation of the isolated ligand (**Fig. 5c**, blue curve) and ligand refinement within the protein’s environment (**Fig. 5c**, green curve). The ligand-only refinement results in stronger decreases in deformation energy, reflecting over-relaxation of the ligand’s structure without the protein structural constraints. Fitting the deformation energy decay as a function of the number of IPA iterations using power-law functions indicates an expected deformation energy of approximately 35.0 kcal.mol^−1^ after ten cycles, when including the protein environment, compared to approximately 15.0 kcal.mol^−1^ for the ligand-only refinement. This highlights the importance of the protein environment in achieving chemically realistic ligand representations within the binding site for highly distorted structures. Re-running the refinement pipelines led to minor improvements in RSCC.

Another example is provided by the PDB entry 3zy2 (CCD-ID: GDP)^24^ (**Fig. 5d**), which contains two alternate conformations of guanosine diphosphate (GDP) bound to the glycosyltransferase POFUT1. Adjacent to GDP’s pyrophosphate moiety, a Mn^2+^ ion coordinates a water molecule. Both alternate ligand conformations exhibit substantial distortions (**Fig. 5e**), with calculated distortion energies of 712.1 kcal.mol^−1^ and 585.4 kcal.mol^−1^, accounting for the atypical IPA overstabilisation of GDP in the protein’s environment (**Fig. 5f,g**).

To assess the role of Mn^2+^ in refinement, two three-step IPA curation strategies were followed: ligand-only refinement and refinement including Mn^2+^ and its coordinated water molecule (**Fig. 5g,h**). Although both approaches reduce ligand distortion, excluding the metal ion led to a greater structural drift, disrupting the interaction networks of pyrophosphate and Mn^2+^ (*e.g.*, with R40 and K71). Including the metal ion preserves these interaction networks, consistent with the stabilising role of cations in highly anionic groups, like the pyrophosphate.

The effect of iterative refinements was quantified by plotting IPA binding energies as a function of refinement cycles (**Fig. 5f**). As previously observed, the largest distortion correction occurs in the first iteration (IP-L-i1), with progressively minor refinements in subsequent cycles. The final IPA binding energies of the two ligand-only optimisations converge to identical values, reflecting the structural degeneracy proposed in the original model. Incorporating Mn^2+^ during quantum mechanical refinement (IP-L-Mn-i3) differentiates the total binding energies: conformer A becomes more stabilised relative to the ligand-only refinement (IP-L-i3), while conformer B was destabilised. This suggests that the PDB-deposited conformational degeneracy may result from crystallisation artefacts. The effect of H238 protonation was likewise investigated, as default algorithms typically protonate this residue at the δ-position. IPA shows that an ε-protonation is more reasonable, as this tautomeric switch lowered the system’s energy in all analyses (**Fig. 5d**).

Finally, IPA’s fragment analysis mode was used to calculate the binding energies of GDP in complex with the most relevant peptide, metal ion, and water components (**Fig. 5i** and **S11**). Across multiple states defined by residue orientations (**Fig. 5j**), the lowest binding energies are consistently obtained when Mn^2+^ is included in ligand refinement, supporting the relevance of chemically consistent modelling. These calculations further support the reorientation of H238, enabling its Nε atom to become a hydrogen bond donor to GDP’s N7, allowing H238’s Nδ to accept a hydrogen bond from S358.

The refinement pipeline was re-run for the IPA optimised structures, and this model, compared to the original one, improved RSCC by 0.09 units and removed several clashes (**Fig. S11**).

### Resolving transition metal coordination in protein binding sites

We applied IPA to a transition metal-containing system. The PDB 3wml^25^ features two metal centres–Fe^2+^ and Co^2+^–coordinated to a post-translationally modified lysine residue (KCX169). In the PDB-deposited model, both metals are coordinated by Cε histidine atoms (**Fig. 6a**). IPA generated sixteen alternative models by systematically sampling histidine side-chain orientations. Most configurations led to proton-metal overlaps, resulting in structural instability (**Fig. S12**). Expectedly, only the configuration featuring four N-coordinated His residues remained reasonable after IPA optimisation (**Fig. 6b** and **S12a**).

**Figure 6:**
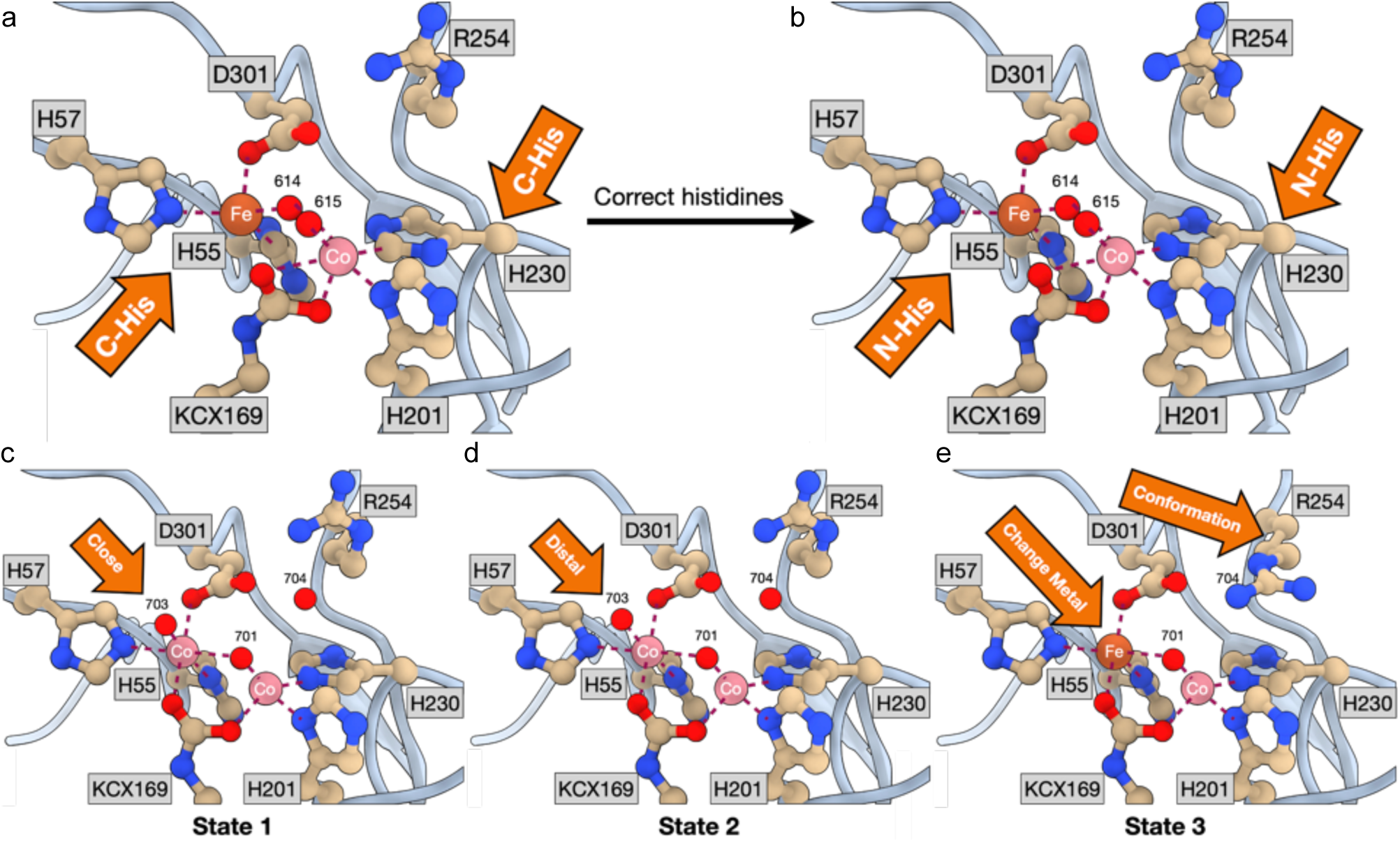
Analysis of the 3wml complex containing two transition metals. a) The PDB-deposited structure of 3wml with two chemically unrealistic C-coordinated histidines. b) After running the IPA protocol, only one conformational state remains, where all metal-coordinated histidine residues do so using either Nε or Nδ. This state avoids metal-proton clashes or the formation of carbenes. The IPA-curated structure was reintroduced in the refinement protocol, from which c-e) three overlapping states were identified. Two states likely contain two cobalt(II) ions, with a bridging water molecule. The two states differ in how water 703 interacts with one of the cobalt centres: c) the water is in the hydroxyl form and coordinated to the metal; d) the water is neutral and uncoordinated. e) The third state likely mixes iron and cobalt. The bridging water remains; however, the iron centre has no equivalent to water 703. The conformation and protonation states of R254 are also changed.

Following additional model building to address regions of positive difference density,^26^ the refined structure was reintroduced into the crystallographic refinement pipeline, yielding three distinct overlapping states (**Fig. 6c-e**). Two of these states likely contain two Co^2+^ ions and no Fe^2+^, a metal-bridged water molecule (HOH701), which, according to IPA, is identified as hydroxylated. The distinction between these two states lies in how water HOH703 interacts with the leftmost cobalt ion: in one state, HOH703 coordinates directly to the metal and is also hydroxylated (**Fig. 6c**), whereas in the other, the Co–O distance increases, and IPA identifies HOH703 as an uncoordinated neutral water molecule (**Fig. 6d**). The third state (**Fig. 6e**) features a mixed-metal configuration with one Fe^2+^ and one Co^2+^ ion, with residue R254 changing its conformational state. The position of H201 favours a neutral form of R254 to avoid steric clashes involving the Hε atom. **Fig. S13** shows the optimised structures and their calculated relative energies.

## DISCUSSION

Structural biology has undergone transformative advances in recent years, driven by breakthrough algorithms built on decades of community effort and the foundational infrastructure of the PDB. Yet, experimental structure-determination methods remain fundamentally limited by resolution: many atoms are not directly observed, and atomic identities are often not unambiguously assigned. To address these challenges, validation tools have been developed to assess structures against established chemical principles. However, accurately resolving the chemistry within macromolecular binding pockets remained elusive. Misassigned atomic identities lead to flawed interpretations–by both humans and algorithms–with broad downstream consequences for drug design, functional annotation, application of machine learning (ML), and beyond. Building on our work with protein-ligand databases,^9^ we estimate that up to 30% of structural models may be inaccurate. Our tailored QM analysis introduces chemical heuristics to augment experimental data, thereby improving model fitness and revealing errors that conventional methods would otherwise miss. Most importantly, IPA’s heuristics extend beyond chemical applications and demonstrate the real impact of structural interpretation in domains such as drug discovery.

The main focus of this work was the application and validation of IPA using crystallographic data, supported by the underlying experimental data available from the PDB. However, IPA requires only a structural model as input, which can originate from several sources. For example, in the Supplement S14, IPA is applied to an NMR-derived model, yielding additional refinement beyond the original structure. This allowed, e.g., fixing N-O bonds that were overly distorted in the initial model. The method can equally be used to evaluate and prioritise AI-predicted models. For instance: i) in the context of ligand validation, as illustrated with the curcumin example discussed above; ii) by identifying the model that exhibits the most comprehensive interaction network, as further explored in the Supplement S15; iii) by refining selected sets of bond distances within a model, as shown in Supplement S16; iv) by guiding selectivity. Crucially, owing to its efficient partitioning of the biological system, IPA scales effectively with system size. This is exemplified in the analysis of cryo-EM models in Supplement S5, where the deposited PDB structure comprises nearly 73,000 atoms, or in the analysis of Protein-Protein Interactions, as exemplified in S17.

IPA enables chemically informed structural model validation. By assessing the plausibility of local configurations, it identifies the most likely chemical states within biological assemblies. Integrating IPA into structure determination pipelines can enhance model fidelity–retrospectively, by correcting existing PDB entries, and prospectively, by supporting real-time model assessment–and expand into impactful opportunities in emerging disciplines such as quantum crystallography. Furthermore, IPA curation is expected to improve the training of structure-aware protein models by preventing the propagation of chemically implausible configurations that can exacerbate known limitations in predictive accuracy.

## METHODS

### Theory

QM-based methods have previously been used to enhance the accuracy of protein-ligand structural models. Merz and colleagues introduced a QM/MM refinement protocol integrated into crystallographic workflows.^27^ Perola and Charifson,^28^ and later Merz and Stewart,^29–31^ developed biased methods that constrain energy surfaces with harmonic potentials to align with experimental geometries. Here, we leverage instead an experimentally biased QM framework to determine the most chemically relevant ligand states–protonation states and tautomers–within the environment of a biological macromolecule. This is achieved by restraining the molecular energy surface onto an experimental structure using a metadynamics Gaussian potential:^32^

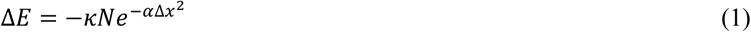

Δ*x* is the root mean square deviation (RMSD) towards the reference experimental structure, which is the target molecular conformation. *κ* and *α* are parameters in the model potential. We consider only positive *κ* and *α*, such that the bias potential Δ*E* is attractive towards the reference structure. *N* is a factor we take as the total number of atoms used to calculate the RMSD. This setup allows constrained minimisation while exploiting the efficiency of unconstrained optimisation algorithms. Ideally, *κ* should be in the range of 15-30 kcal.mol^−1^, whereas *α* should take values within 0.5 and 1.0 Å^-2^. This allows rapid, systematic optimisation of structures while retaining conformation: bond distances and angles are the primary variables optimised. This is key to reducing the distortion caused by the limited resolution of experimental techniques.

### Using IPA

IPA is available in two scripts. The main script, *In-Pocket-Analysis.py*, is summarised in **Figure S18**. Briefly, the central residue (ligand) is extracted from a PDB or mmCIF file. By default, all residues within 5.0 Å are considered for the calculation. The structures may be protonated using OpenBabel,^33^ RDKit,^34^ or a chemical component dictionary (CCD) downloaded from the PDB. The CCD-protonation uses ULYSSES’ internal libraries. The search for ligand protonation and tautomeric states is performed with RDKit. The latter is also used to include various amino acid-protonation states. RDKit considers two protonation states for aspartic and glutamic acids, lysines, and tyrosines. For histidines, four protonation states are considered. After each run, an SDF file for the ligand is generated, which can be used to restart calculations. By default, each amino acid is N- and C-capped. We use the closest carbonyl group coordinates and add a hydrogen atom to form an aldehyde (C-terminal) or formamide (N-terminal). The capping effect is also subtracted from interaction energies by default. The program’s output includes a text file containing general information (deformation energy, distortion energy, summed interaction energy, relative tautomer energy–if several tautomers are considered, tautomer’s total energy, and RMSDs), the ligand-residue interaction energies in a CSV file, all optimised structures, and the outputs of each QM calculation.

The default RDKit run lists all possible chemical states available to the system and found by the algorithm. In some situations, this includes chemical states with negligible biological and chemical relevance, like imidic acid variants of axitinib (CS2 and CS4 in **Fig. 2b**). When running IPA, the user may run calculations on all chemical states or, if prior chemical knowledge is available, specific, preferred ones.

### Performance

For a ligand of the size of axitinib with ten chemical states, considering a radius of 4 Å around the central residue (*ca.* 14 residues and their additional protonation states and flips), IPA takes about 3 minutes on an Apple M2 Pro laptop.

### Availability

IPA is available from the ULYSSES library.^35^ A stand-alone tool will be available in https://github.com/s1riius/In-Pocket-Analysis.git upon publication. The IPA code will be freely available for academic users and applications. IPA is already available on EMBL Grenoble’s CRIMS platform,^36^ and we are currently integrating IPA into CCP4^37^ and PDB-REDO.^20^ Note that, due to the specificities of some licenses, not all protonation engines might be available from all interfaces.

### Limitations

The primary limitation of IPA lies in its treatment of only pairwise (residue-residue) interactions, which often fail to capture the full complexity of biological environments. Notably, this includes regions with localised accumulation of carboxylate groups or coordination spheres involving transition metals–situations where higher-order interactions and cooperativity effects are critical. To address such cases, we introduce the IPA-fragment mode, which was essential in several parts of this study. This extension enables the treatment of more chemically complex environments, though at a higher computational cost: the scaling increases approximately cubically with system size. Users are therefore advised to consider this when setting up systems for further calculations.

Finally, while IPA-derived binding energies offer valuable insight into the chemistry of binding pockets, caution is warranted when applying them as scoring functions, for instance, in drug discovery workflows. Although such use is briefly explored in this work, broader validation is needed before routine application in this context. We recommend using binding energies from the fragment mode.

### Data Availability

All the IPA-optimised ligands from the PDBbind^8^/MISATO^9^ databases are available in https://gitlab.com/siriius/ipa-ligands.git. Here, QM-refined geometries and the respective molecular properties are available for download.

## ACKNOWLEDGEMENTS

This work was supported by the Bundesministerium für Bildung und Forschung (BMBF), projects SUPREME (number: 031L0268) and DATIPilot Sprint – MISATO (03DPS1234); by the Bundesministerium für Wirtschaft und Klimaschutz (BMWK) via ZIM (grant number: KK5710001BA4); by the Deutsche Forschungsgemeinschaft, DFG grant CRC387, project ID TRR 387/1-514894665.

## SUPPORTING INFORMATION

**Figure S1:**
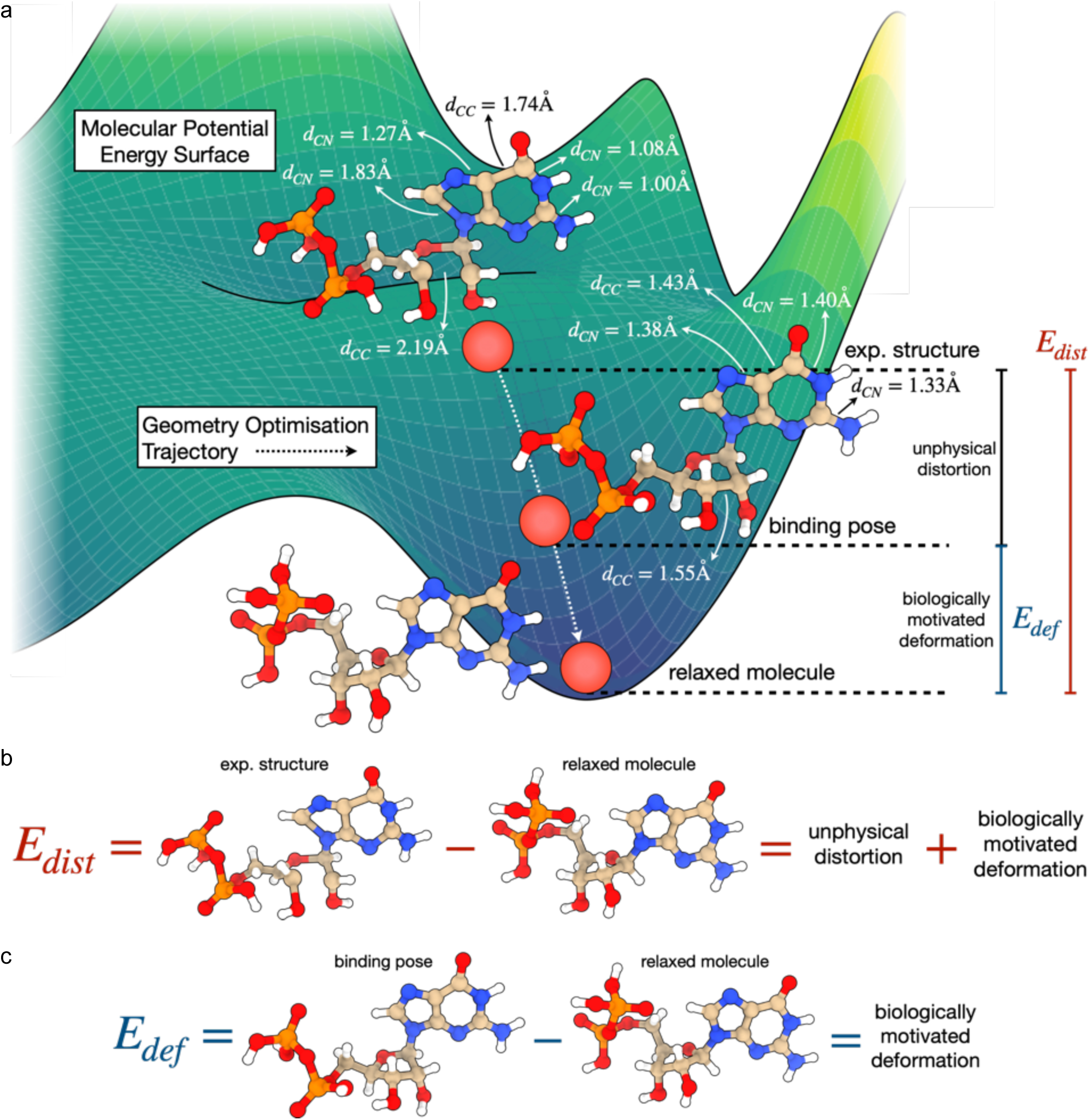
Theoretical background depicting the rationale behind IPA and how distortion and deformation correlate with one another. a) Schematic representation of the various ligand conformational states on the ligand-only potential energy surface (PES). The experimentally observed ligand structure (“exp. structure”) is depicted as the red sphere at the top, corresponding to the molecular geometry shown above it. This structure includes biologically meaningful distortions induced by the biomolecular target, as well as unphysical distortions resulting from limited experimental resolution and modelling inaccuracies (*e.g.*, incorrectly applying restraints while solving a structure). Note that unphysical distortion may arise for lower resolution structures because then the structural model becomes heavily reliant on correctly using and the validity of distance and angular restraints. Examples of such inaccuracies include abnormal bond lengths and angles present in the PDB-deposited model, but can also include atomic clashes, *e.g.*, by refining structures in the absence of hydrogen atoms. In this illustration, we imagine performing a structural relaxation of the ligand in the absence of the protein. The geometry optimisation trajectory is assumed to initially correct the inaccurate bond lengths and angles while maintaining the ligand’s binding pose conformation. This stage primarily removes unphysical distortions from the structure. The trajectory then proceeds to fully refine the ligand’s geometry, relaxing torsional angles and, therefore, also the binding pose. Ultimately, this yields a representation of the free ligand in solution, absent from the influence of the biomolecular environment. b) Naïve calculation of deformation energies. This approach uses the processed experimental structure from the PDB model and compares its energy to that of the fully relaxed ligand, in this case, QM optimised. As illustrated in a), this energy difference includes not only the biologically relevant deformation but also unphysical distortions. We refer to this energy as the distortion energy (E_dist_). c) Biologically meaningful calculation of deformation energies (E_def_), *i.e.*, including exclusively the protein-induced change in ligand conformation. This method uses chemically reasonable representations of the ligand–its binding poses without unphysical distortions in bond lengths and angles–and compares them to the fully relaxed, quantum mechanically optimised structure. In this work, we introduce IPA optimisation, a method capable of refining ligand structures while preserving their initial binding conformations. This enables structural relaxation in the absence of the protein binding site. According to our definitions, E_dist_ encompasses E_def_–the biologically meaningful deformation energy–along with additional, unwanted structural distortions resulting from limited resolution of the experimental data. While one could define E_dist_ to include only these unphysical distortions, doing so would not reflect how deformation energies are typically calculated in practice, nor would it allow for a meaningful comparison between E_dist_ and E_def_. The deformation energy is an important quantity to estimate to study biomolecular binding, as this quantity has a negative impact on binding thermodynamics. If ligand deformation energies are unrealistically too large, binding cannot take place. It is therefore important to correctly estimate E_def_ in order to obtain a more realistic representation of a protein-ligand complex.

**Figure S2:**
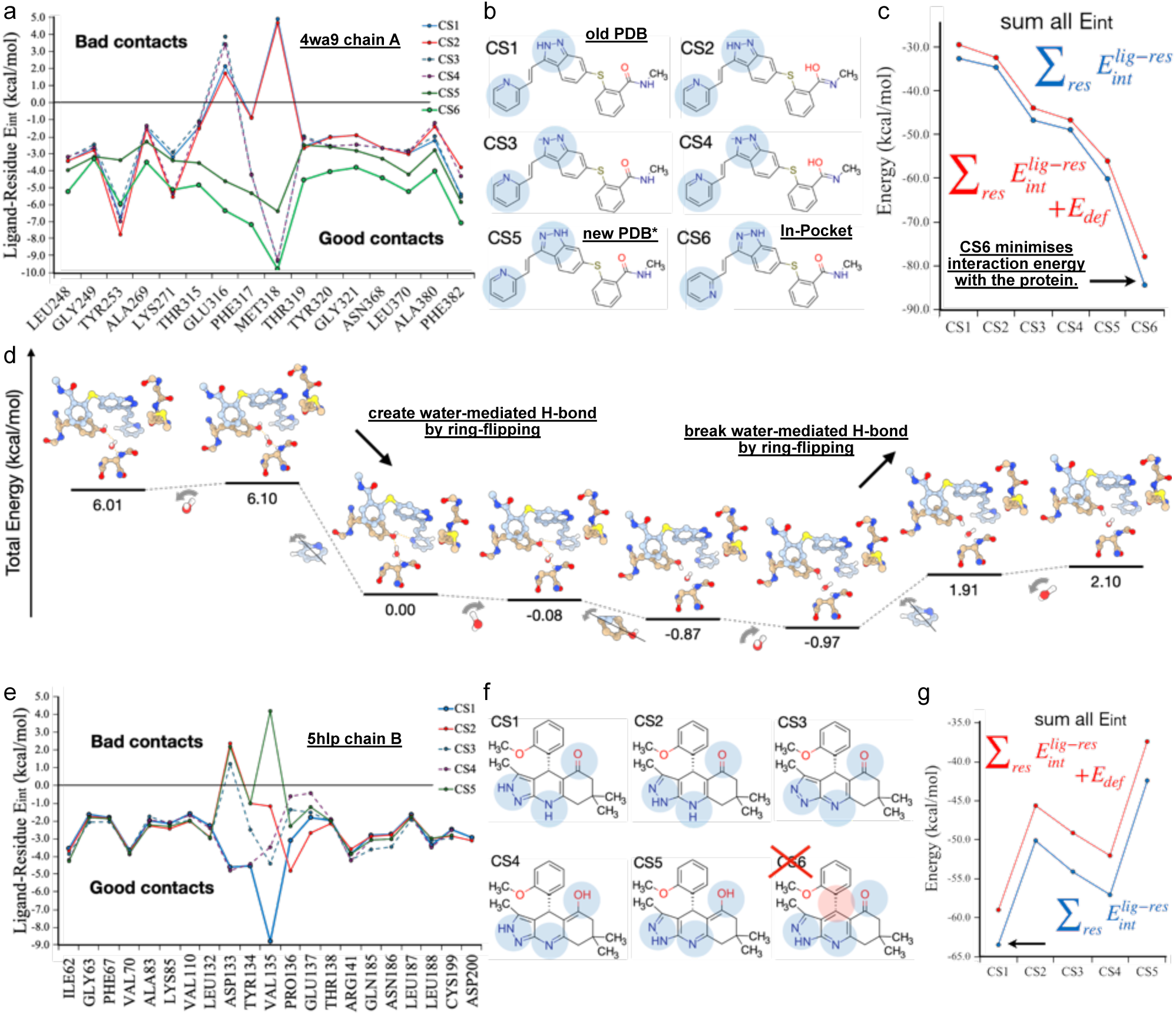
IPA data for other chains of PDB-IDs 4wa9 (chain A, blocks a-d) and 5hlp (chain B, blocks e-g). a) The ligand-residue interaction energy plot shows that CS5 and CS6 are the only two chemical states that do not clash with the kinase’s hinge. Of these two CSs, CS6 offers predominantly better interactions with Y253 and M318. Note that although CS5 and CS6 differ only by a 180° flip of the terminal pyridyl, this leads to slight differences in the IPA refined ligand structures. These fine differences justify the observed shift between the CS5 and CS6 curves. Interestingly, the energy gain by flipping the terminal pyridyl from CS5 to CS6 regarding the interaction with Y253 is not as pronounced as for chain B. b) The chemical states identified by IPA, with structural changes highlighted. c) The summed interaction energies and the total IPA binding energy (summed interaction energies and the deformation energy) show that CS6 is still the most stabilised axitinib chemical state in ABL1’s pocket. d) A fragment interaction energy plot showing that the pyridyl flipping stabilisation with Y253 is, in fact, a water-mediated hydrogen bond for chain A. This shows that the enthalpy stabilisation with Y253 is partly transferred to a water-mediated interaction. e) The ligand-residue interaction energy plot for 5hlp, chain B. Despite minor numerical differences, the results fully agree with those for chain A. f) The ligand chemical states identified by IPA, with differences highlighted in blue or red circles. Note that the PDB-deposited topology is inconsistent with the available structural model. g) The summed interaction energies and the IPA binding energy for the different chemical states show that CS1 is the most stable chemical state of this ligand in GSK3β’s pocket. This is followed by CS4. The difference in binding energy also reflects the loss of an H-bond. **Experimental data:** For accessing all the data underlying the results and the experimental validation using traditional crystallographic metrics, check the “supplements/figure2+S1/” folder, followed by the PDB-IDs.

**Figure S3:**
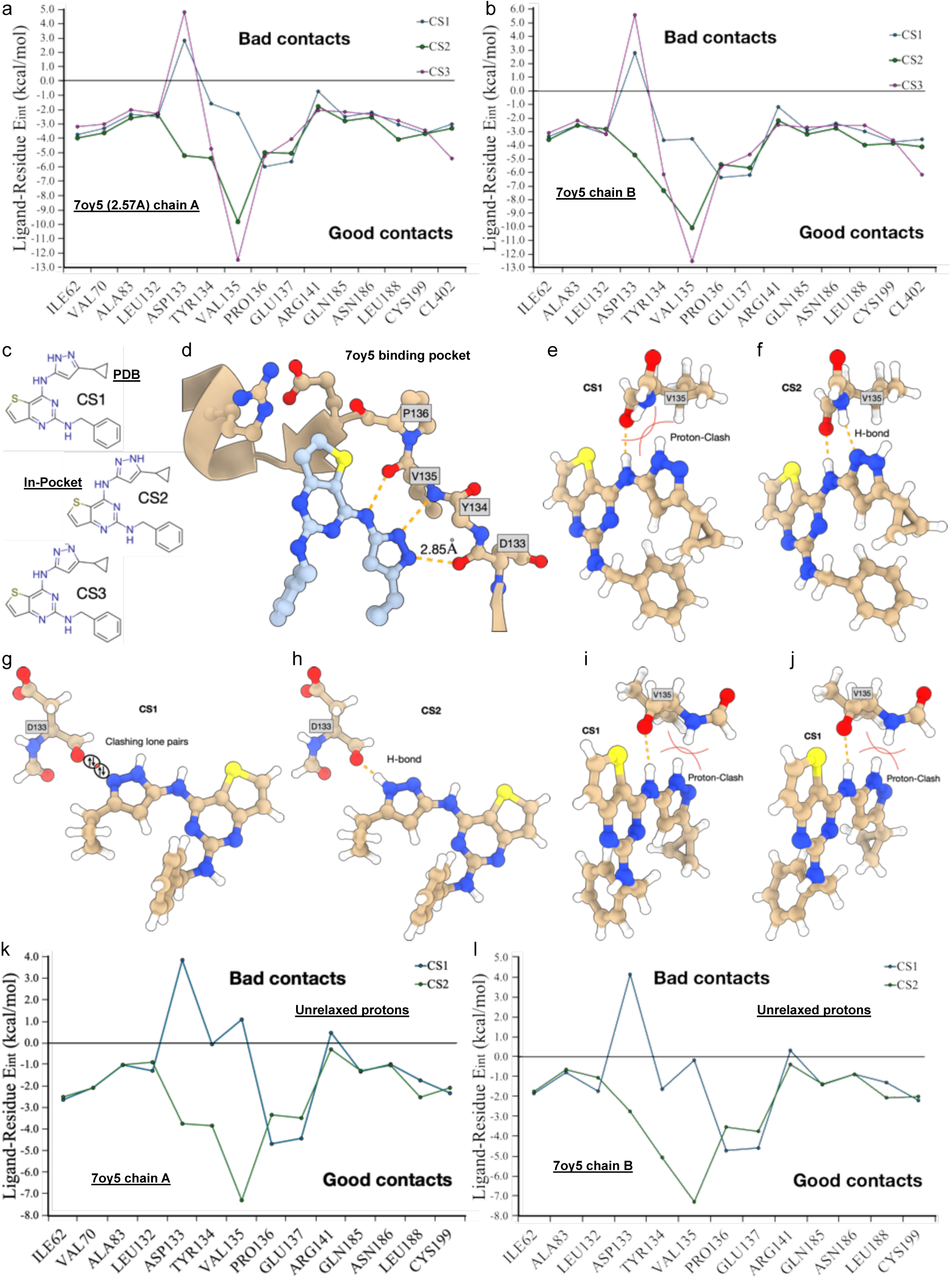
IPA data for a pyrazole-containing ligand bound to GSK3β, data from PDB-ID 7oy5.^38^. a) The ligand-residue interaction energy plot for chain A shows that, of the 3 chemical states identified, only CS2 does not clash with the kinase’s hinge. c) The same plot for chain B. c) The IPA-identified ligand chemical states. CS1, the PDB-deposited topology, differs from the IPA-identified chemical state, CS2. d) Pocket schematic representation with possible interactions from the different chemical states. e) The IPA-optimised CS1-V135 pair shows a proton clash with the main chain amide, avoided during the geometry optimisation by changing the hybridisation of the amide’s nitrogen. f) The CS2-V135 pair shows the hydrogen bond missed by the PDB-deposited topology. g) The CS1-D133 pair shows a clash of lone pairs. Unlike a proton clash, the lone pairs cannot avoid one another. h) The CS2-D133 interaction shows the formation of a hydrogen bond. i) The chain A and j) chain B representations for the CS1-V135 optimised pair with all protons included in the bias potential. The protons cannot avoid each other in this situation, and the clash is more prominent. The ligand-residue interaction energy plot for k) chain A, and l) chain B, when including protons in the bias potential. Here, only minor refinements from the initial structures are possible, and the clashes are more evident.

**Figure S4:**
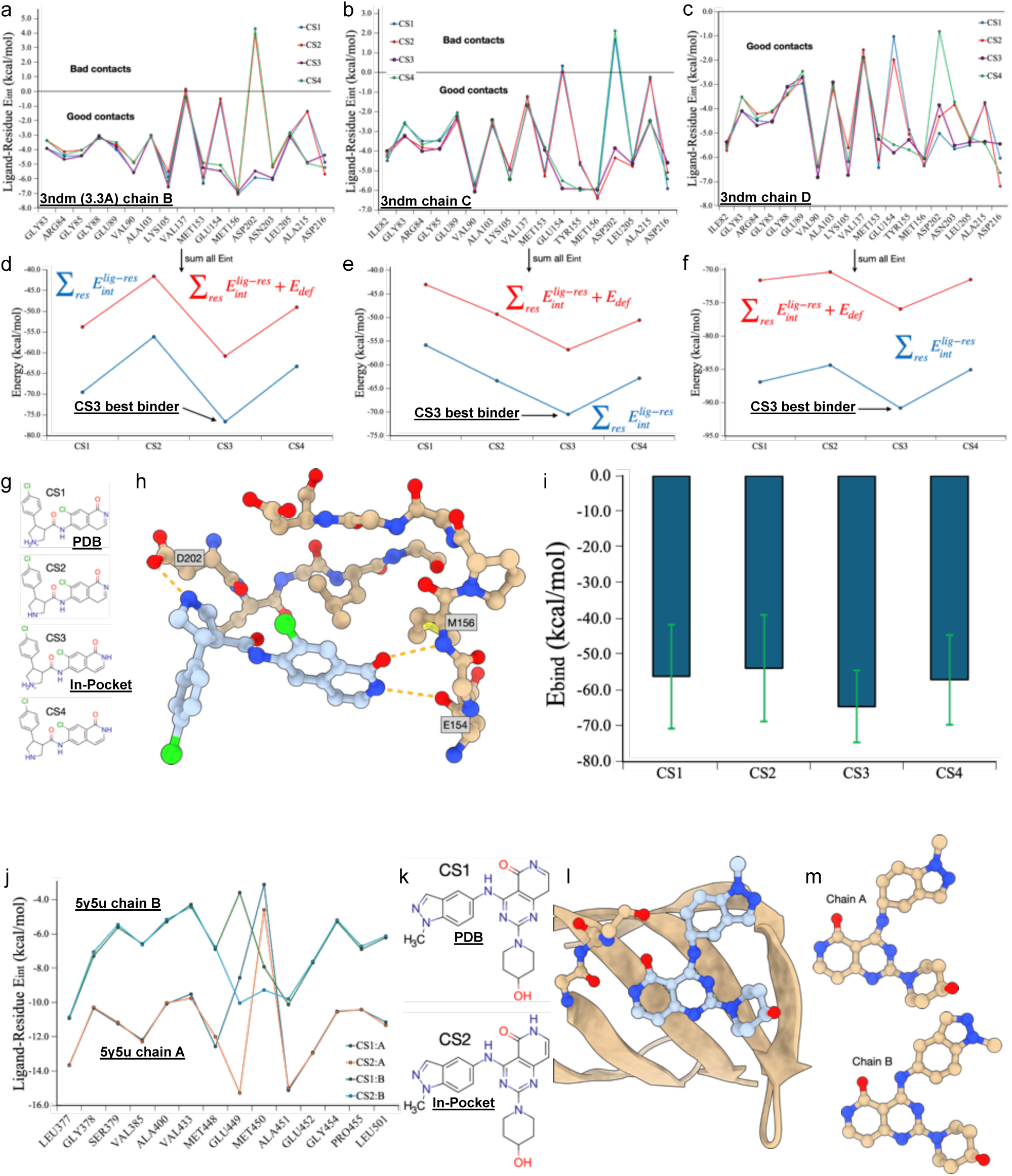
IPA data for isoquinolone-lactam equilibria, data from PDB-IDs 3ndm (a-i)^39^ and 5y5u (j-m). ^40^. The ligand-residue interaction energy plot for chains a) B, b) C, and c) D of the 3ndm model. Note that no ligand was identified in chain A. The summed interaction energies (blue) and the IPA binding energies (red) for chains d) B, e) C, and f) D show that the lactam CS3 offers a better representation of the ligand in the pocket of the ROCK1 kinase. This contrasts with the PDB-deposited topology CS1. g) The IPA-identified ligand chemical states. h) ROCK1’s pocket representation with the 3ND ligand. i) The statistical analysis of the IPA binding energies shows that, within the variability of one PDB file (3 chains), CS3 is clearly the best binding conformational mode. j) The ligand-residue energy plots for another isoquinolone-lactam equilibrium, 5y5u. Both chains are represented in one plot. There is a remarkable difference in the range of stabilising interactions for each chain. k) The ligand chemical states identified by IPA. l) Tyrosine kinase’s pocket representation with the chain A ligand. A significant distortion in the central linker is observed. m) Comparing the ligand structures in the two chains of the PDB 5y5u shows that chain A is unreliable. Analysing all the data, it is clear that although the isoquinolone CS1 does not induce any clash, it lacks hydrogen bond interactions with E449 and M450.

**Figure S5:**
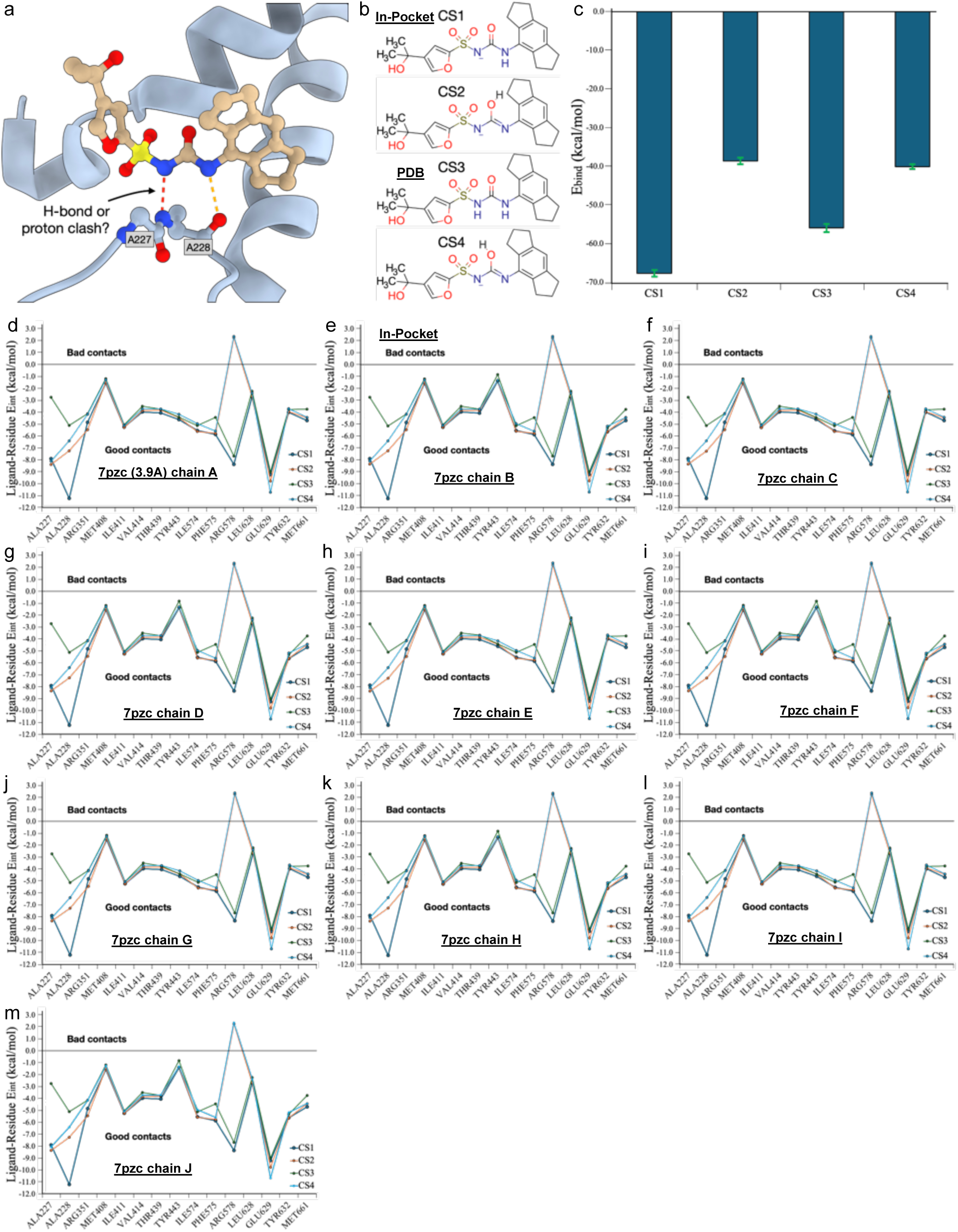
IPA data for the sulfonyl-urea ligand in the PDB 7pzc.^41^. a) The ligand-in-the-pocket representation shows a potential clash between one of the sulfonyl-urea protons and the main chain nitrogen of A228. b) The IPA-identified ligand chemical states. c) IPA binding energy analysis over the 10 chains available in the PDB file. The calculations systematically identify CS1 as the most likely chemical state of the sulfonyl-urea ligand bound to the protein. d-m) The ligand-residue interaction energy plots for each chain. Though proton abstraction from the sulfonyl-urea might initially seem odd, this result greatly agrees with the sulfonyl-urea pK_a_ in the range of 4-6.^42^

**Figure S6:**
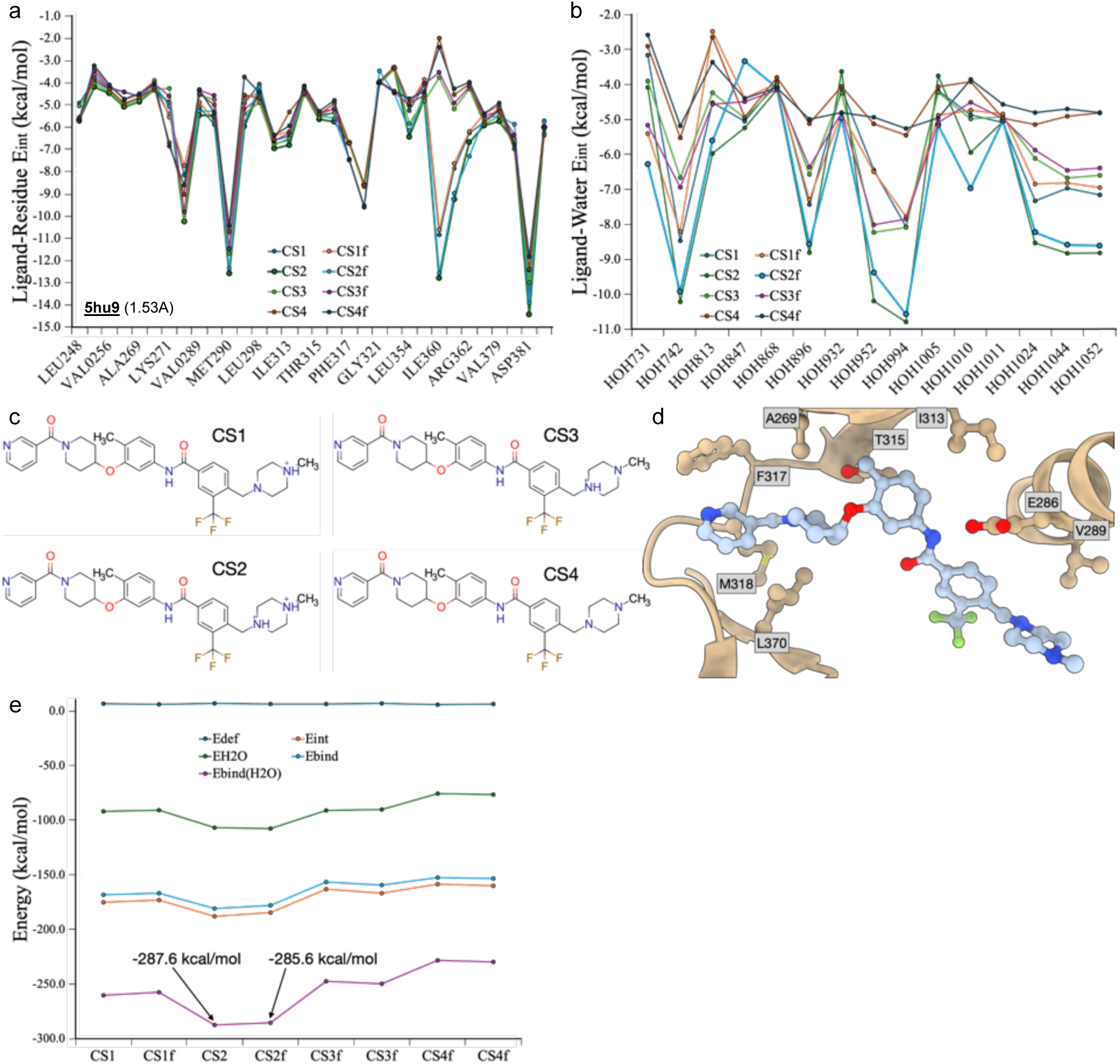
IPA analysis of an N-methylpiperazine ligand with a potential pyridyl flip. a) The ligand-residue interaction energy plot for the protein-ligand complex in the PDB 5hu9.^43^ b) The ligand-water interaction energy plot for the structural model in 5hu9. The analysis was performed for the four IPA-identified chemical states with and without terminal pyridyl flip (suffix “f”). The ligand-residue interaction pattern is similar for all chemical states except around I360. This indicates an advantage for protonating the most external nitrogen in the piperazine ring. Conversely, the ligand-water interactions seem stronger for CS2 for both pyridyl ring orientations. c) The IPA-identified ligand chemical states. d) The pocket model for the complex in 5hu9. e) The summed energy analysis for this complex: The deformation energy (E_def_), the interaction energy with solvation waters (E_H2O_), the summed ligand-residue interaction energies (E_int_), the IPA binding energy (E_bind_ = E_int_ + E_def_) and the IPA binding energy with the water contributions (E_bind(H2O)_ = E_bind_ + E_H2O_). The last quantity differs by 2.0 kcal/mol for the two flips of CS2, indicating that it is likely that both pyridyl ring orientations are biologically stabilised and observed.

**Figure S7:**
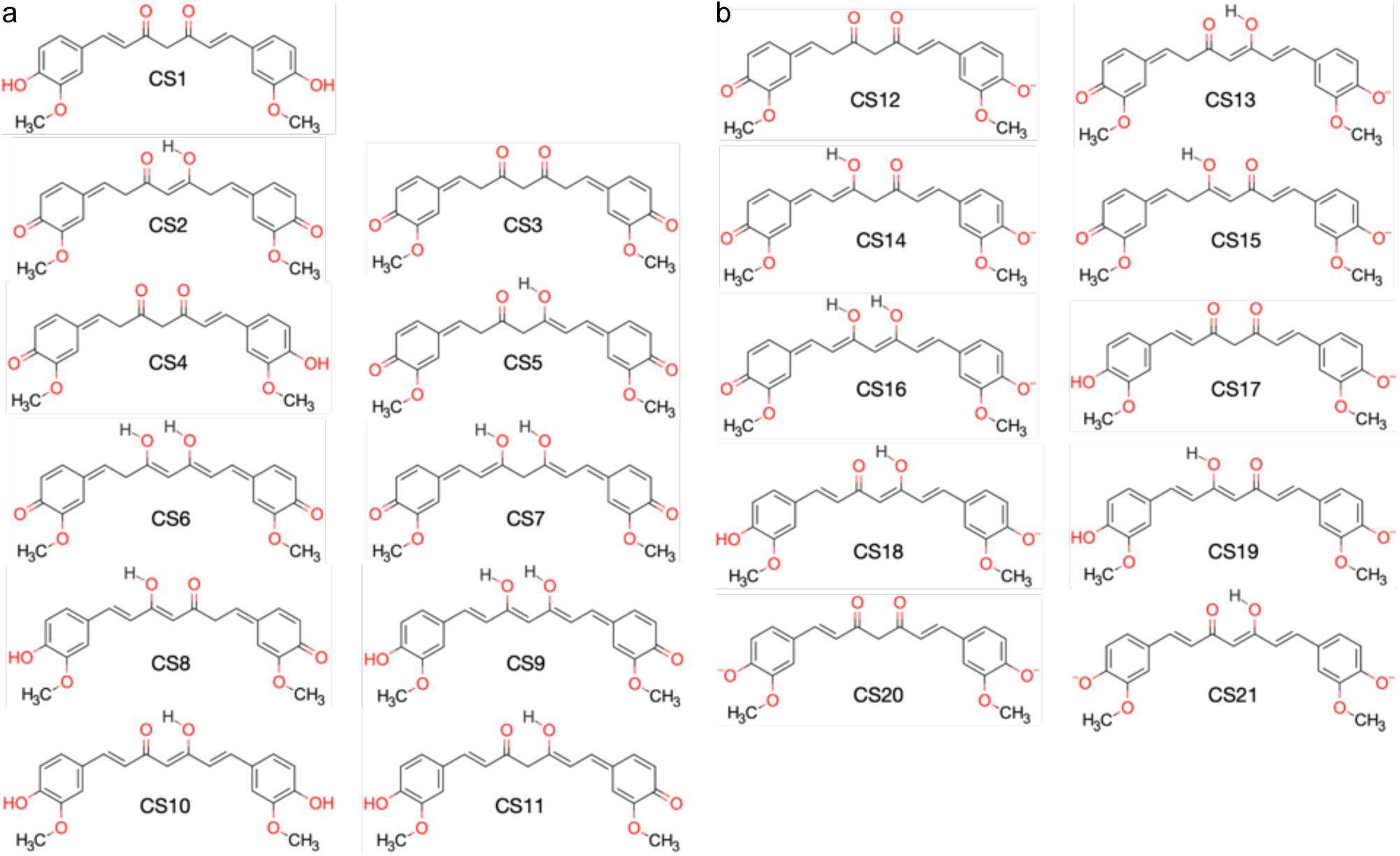
The manyfold curcumin chemical states identified by IPA. a) The neutral chemical states. b) Charged chemical states.

**Figure S8:**
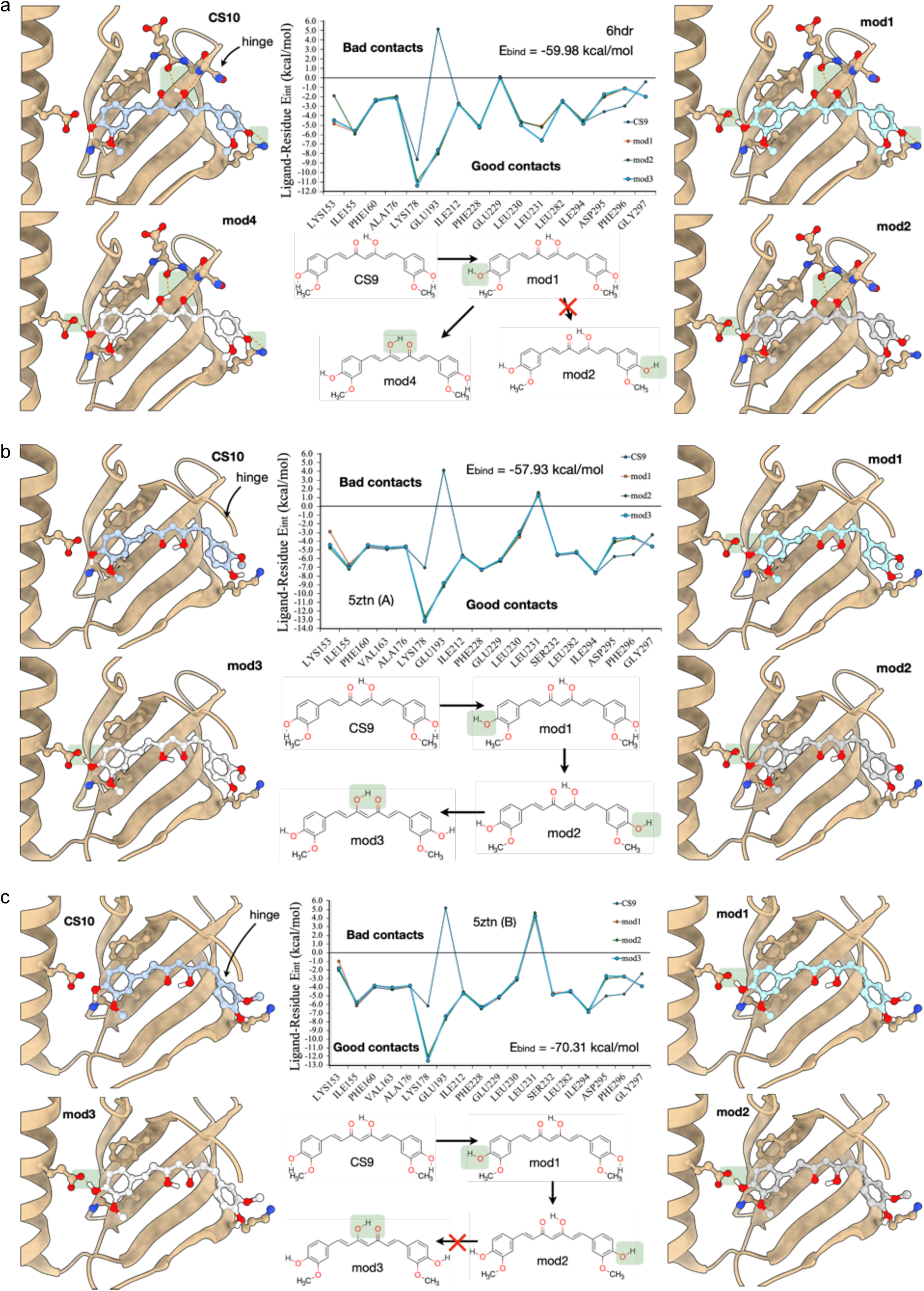
IPA of curcumin with different protonation patterns consistent with curcumin’s CS10 chemical state. The impact of orientation and proton jumps on the protein-ligand complex stability can be followed by the ligand-residue interaction energy plots in the centre of each block. Structural models are placed to the left and right of the interaction energy plots. This data supplements the information in Figure 4 of the main manuscript. a) The hydrogen bond network optimisation for the structural model 6hdr. b) The hydrogen bond network optimisation for the structural model 5ztn, chain A. c) The hydrogen bond network optimisation for the structural model 5ztn, chain B.

**Figure S9:**
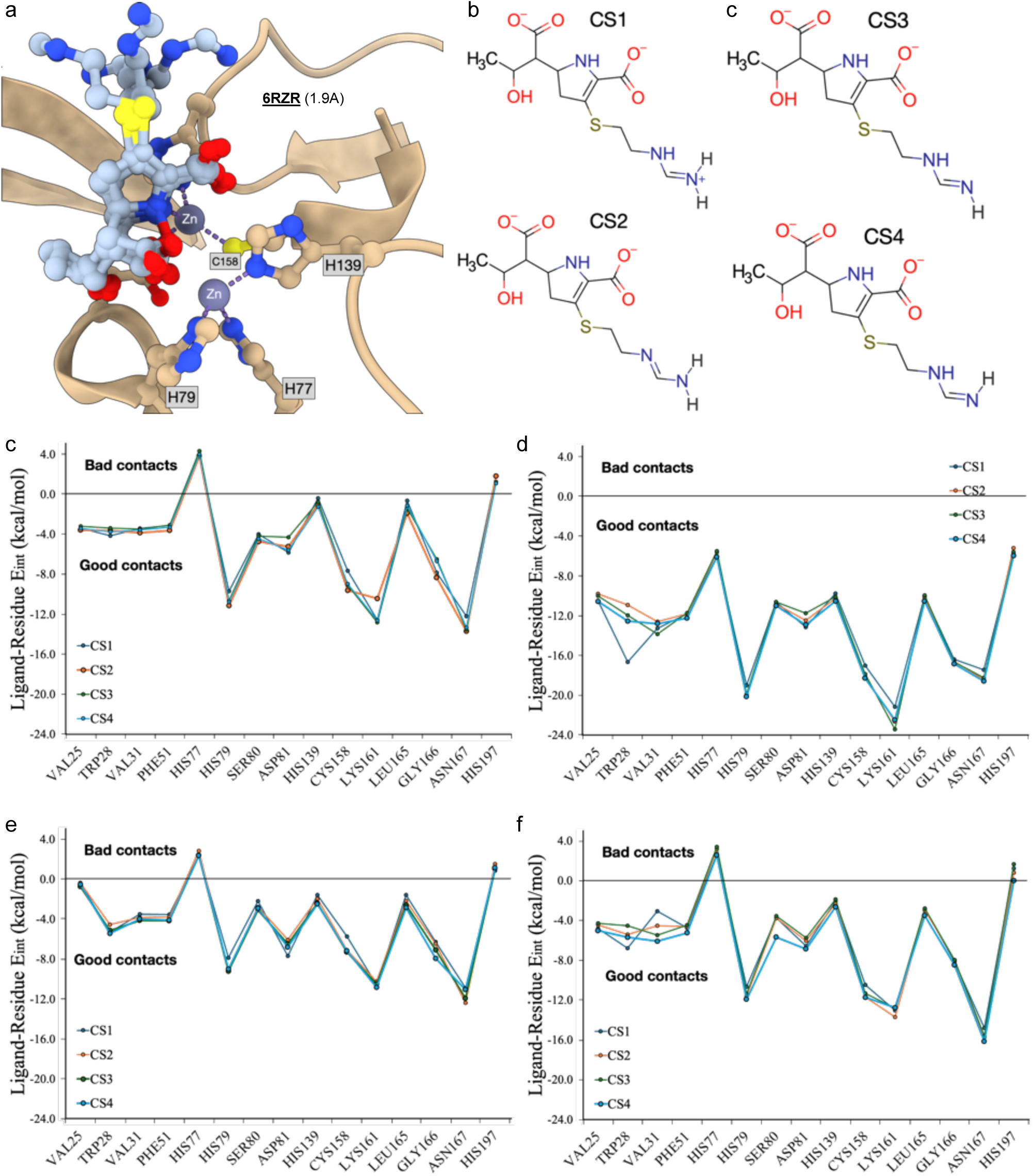
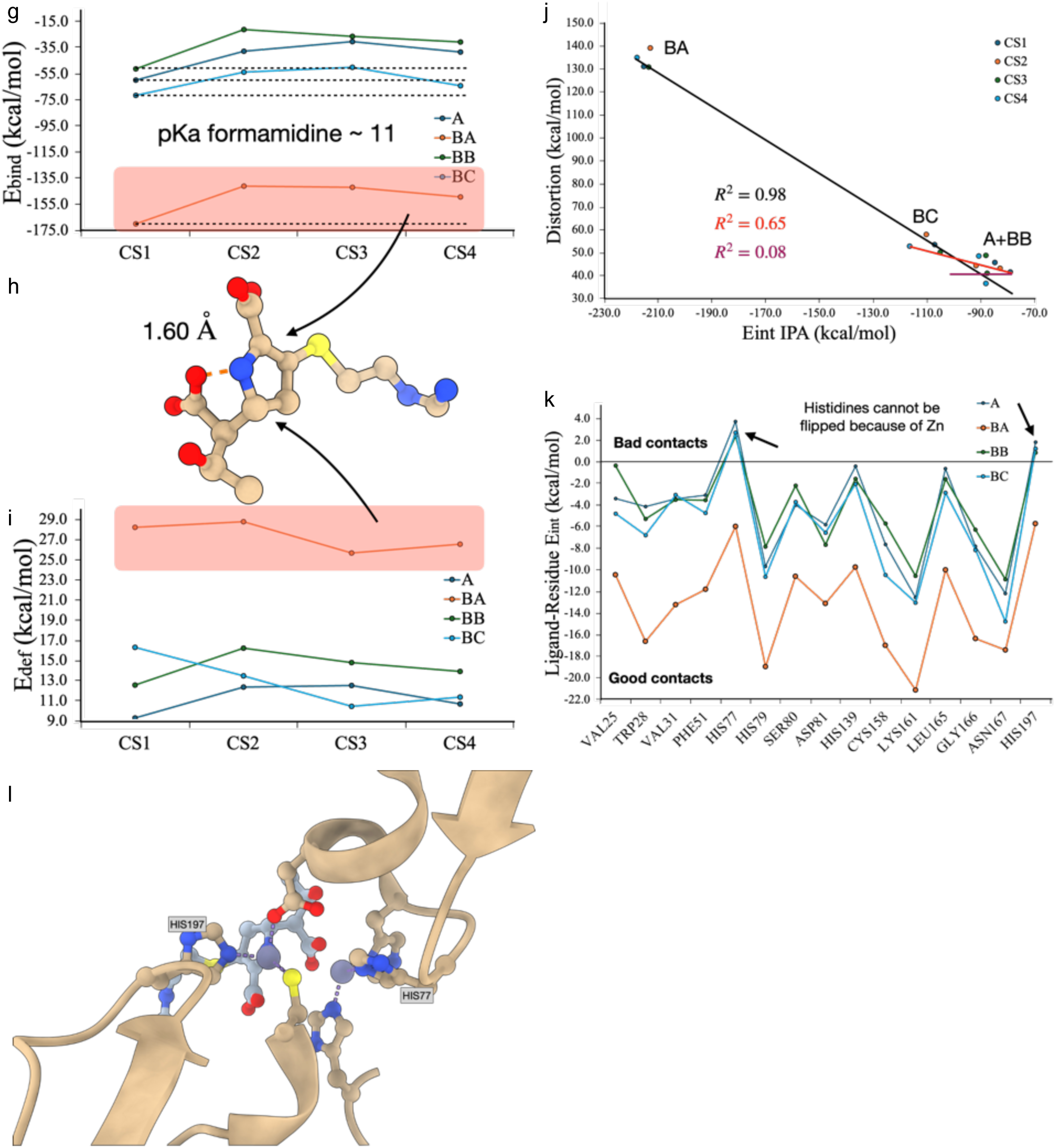
Analysing the alternate locations of a ligand next to a metal-containing binding pocket, PDB-ID 6rzr.^21^. a) The pocket representation, with an overlay of the four structural models: chain A and chain B, the latter with three alternate binding poses (BA, BB, BC). b) The four IPA-identified ligand chemical states available. Note that we purposely excluded the carboxylic acid protonation states. The ligand-residue interaction plots for c) chain A and the chain B models d) BA, e) BB, and f) BC. Although the ligand-residue interaction energy plot for the BA model shows a pattern similar to other models, this is shifted towards lower energies, indicating something odd with this model. g) IPA binding energies also show a different behaviour for the BA model. h) Ligand-only representation of the BA model, evidencing an N-O short contact that is almost within the distance of a covalent bond. This inaccuracy in the structural model raises the i) ligand deformation energies for all chemical states identified. j) The distortion Vs. the IPA-summed ligand-residue interaction energies plot shows a good correlation when the data referring to the structural model BA is included. The correlation is negatively impacted by excluding model BA’s data, but in this case, there is still a strong Pearson correlation between model distortion and the summed IPA interaction energies. This indicates that one of the models is still contaminated with low experimental data resolution. Excluding the BC-model dataset destroys any correlation between distortion and the summed interaction energies, showing that models A and BB are the most reasonable choices for further studies. k) The ligand-residue interaction energy plots for CS1 in several structural models. This shows that, of the two remaining models, chain A is the QM most indicated pocket representation of this complex. IPA shows two poor contacts associated with two histidine residues. l) Analysing the structural model shows that the histidines cannot be flipped; otherwise, the interactions with the zinc ions would be impacted.

**Figure S10:**
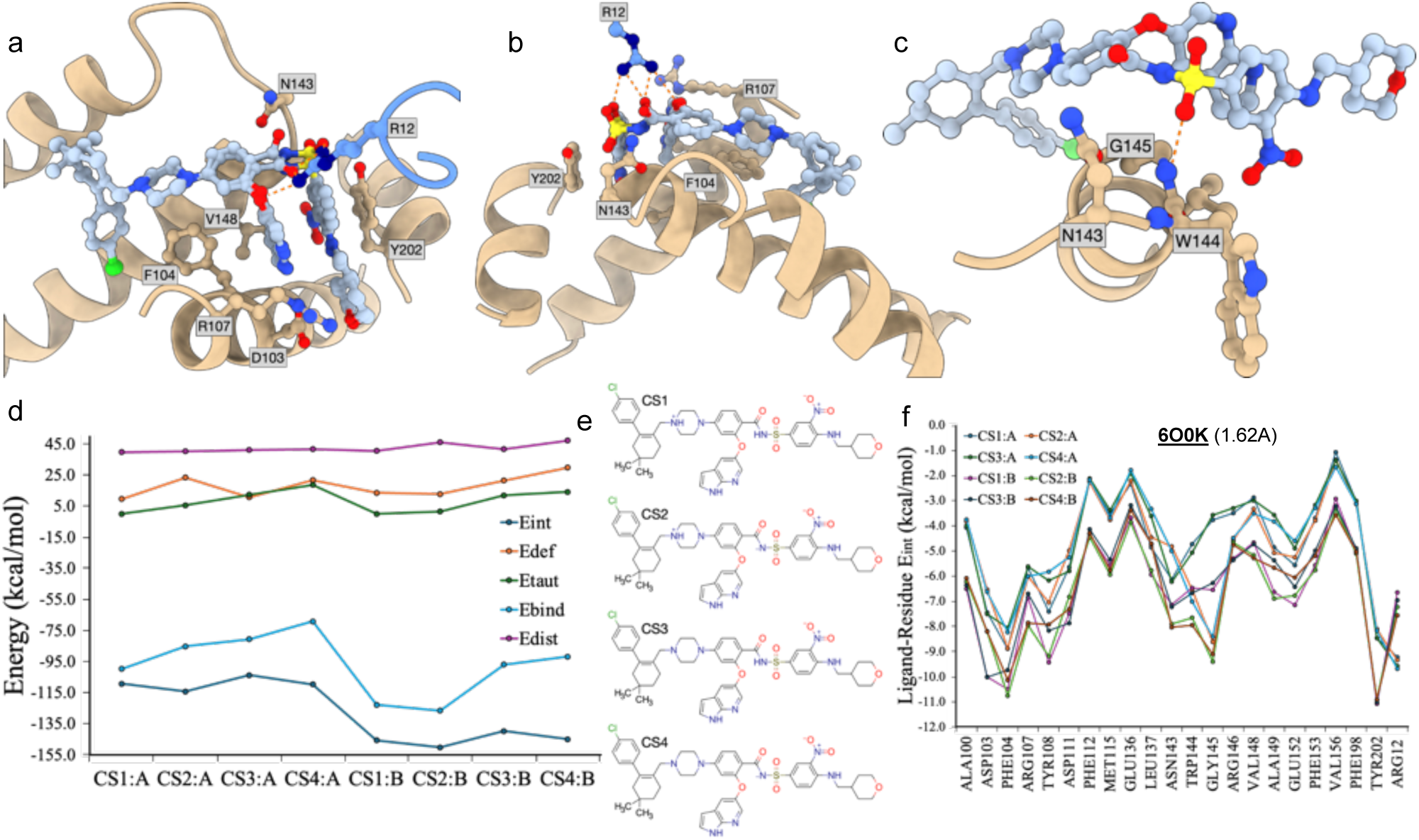
Two venetoclax models in BCL-2, PDB-ID 6o0k. ^22^. a) The pocket of venetoclax in BCL-2 has two alternate ligand representations (A and B). Residue R12 is included from a symmetry copy in the crystal. b) Alternative view/representation of the binding pocket, showing the interaction pattern between venetoclax and the residue R12. c) The interaction of venetoclax with G145. By deprotonating the sulfonyl-amide, charge density is increased in the sulfonyl group, strengthening the hydrogen bond to G145. d) Global IPA metrics for the two structure models. This plot shows that all distortion energies are similar and within the same magnitude. This rules out distortion as the driver for the shift of the interaction energy plot in f), and it leads to the conclusion that alternate B offers a better structural model of venetoclax in the pocket of BCL-2. This is also confirmed by the IPA binding energy, which contains the summed ligand-residue interaction energies, the ligand deformation, tautomerisation, and isomerisation energies. Of the four chemical states investigated, those with the ammonium group are particularly favoured (vs. neutral amine). Deprotonation of the sulfonyl-amide group leads to an additional stabilisation by 4.0kcal/mol, by strengthening the hydrogen bond with G145’s main chain nitrogen. Additional stabilisation is obtained by interactions with water molecules, see supplemental tables and plots in the supplements folder. We stress that one of the main differences between models A and B is the interaction of the ligand with R12. As the target BCL-2 is not a biological dimer, and the interactions with R12 result from symmetry copies, it is likely that this is a crystallographic artefact and that the experimentally observed alternate conformation A is not biologically relevant. e) The four venetoclax chemical states we considered in the analysis. f) The ligand-residue interaction energy plots show a potential shift in the interaction energy pattern of the two alternate models we are analysing, alternate A and alternate B. The almost indistinguishable distortion energies between the two models (see above) show this is not caused by ligand distortion or model quality, but rather differences in the binding pose.

**Figure S11:**
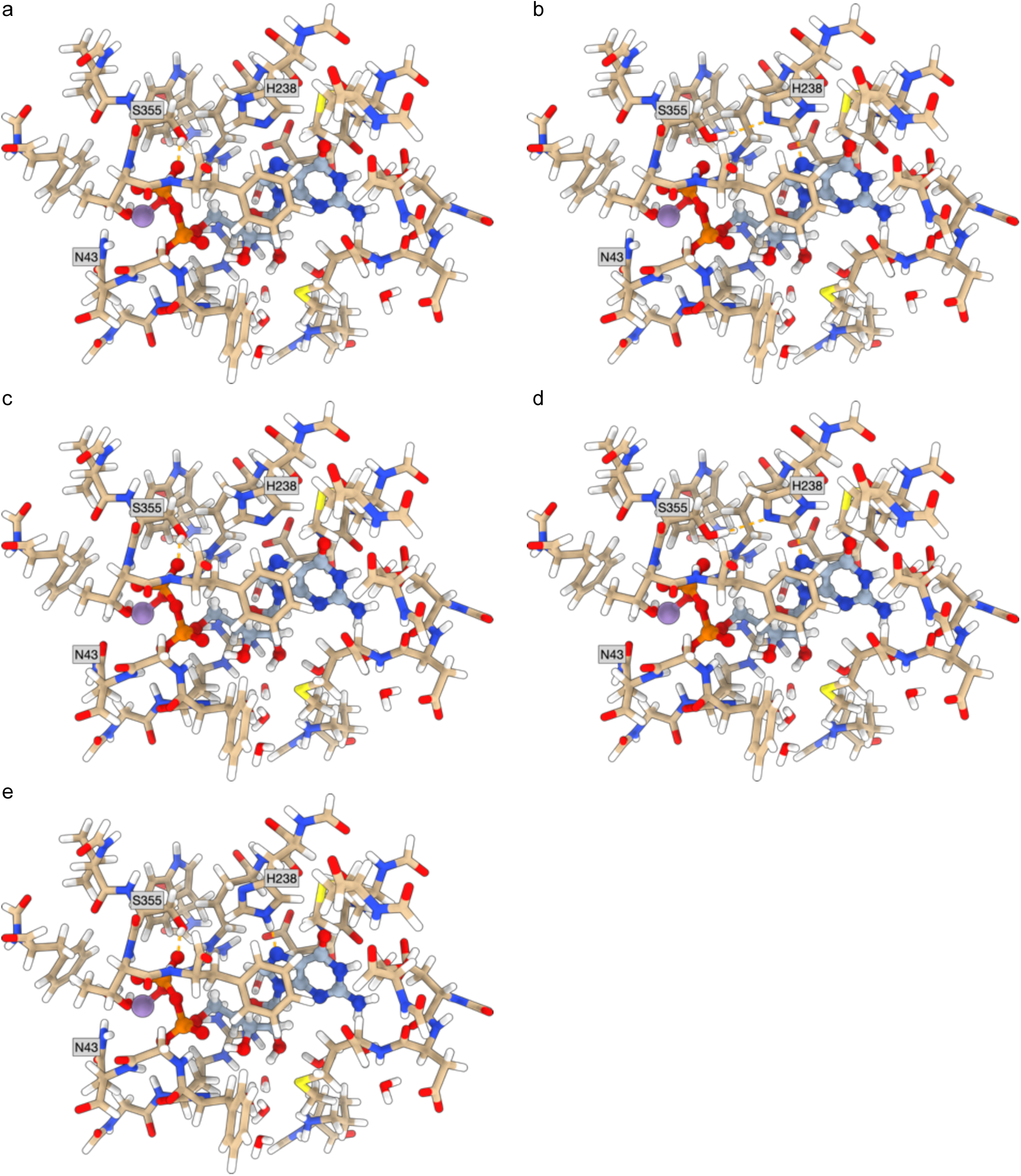
Structural representation of the five 3zy2 conformer states tested. a-e) represent, respectively, the states 1-5 referenced in Figure 5i and **5j** obtained by combinatorially flipping H238 and N43. The last state changes the protonation state of His238 from δ to ε.

**Figure S12:**
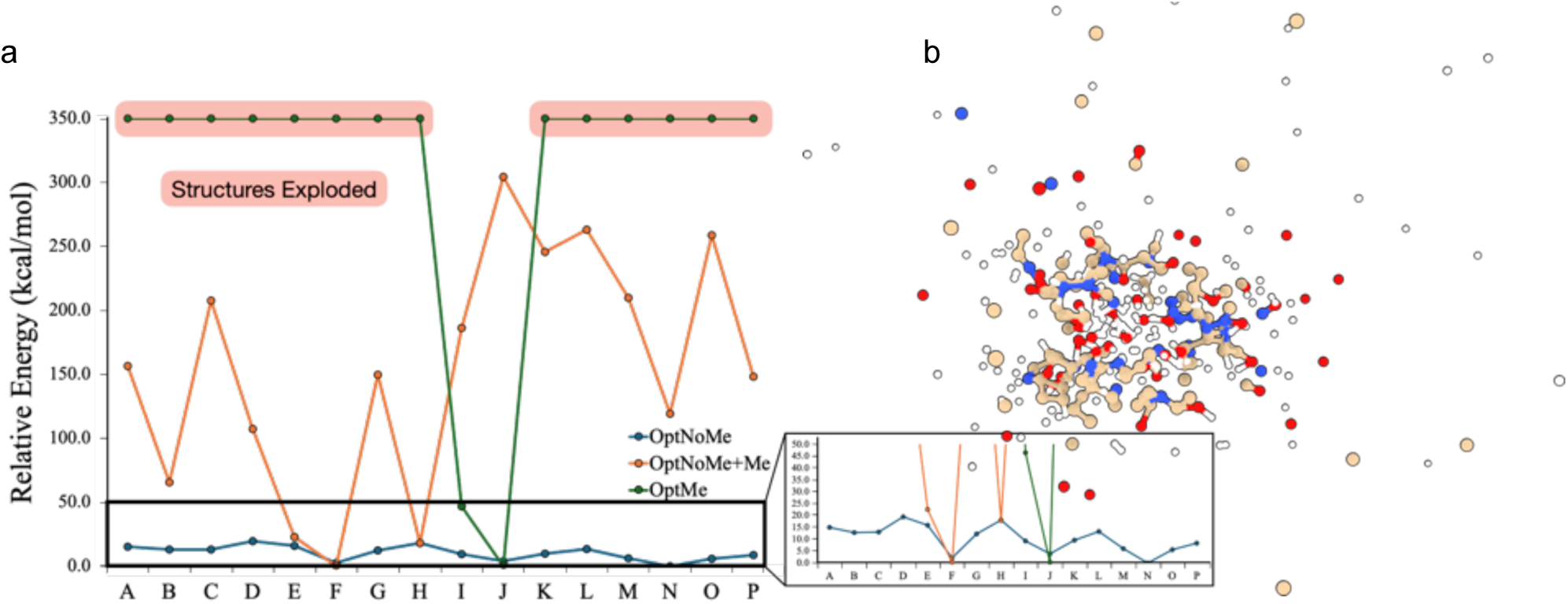
Retrieving chemically meaningful information for large atomic systems. a) The energy plot of all conformational states considered when analysing the PDB-ID 3wml. Three curves are shown: a blue curve, where the fragments were optimised without the metal centres. Though reasonably flat, the energy surface is dominated by a single conformer. Introducing the metal ions on the previously optimised structures, the orange curve severely destabilises several conformers, particularly the expected energy minimum. Finding the energy minimum requires optimising the protein fragments in the presence of metal ions, represented in the green curve. In the presence of metal ions, only two structures can be optimised. Despite the constraining potential used, most structures ended up exploding, for which we pragmatically assigned a relative energy of 350.0 kcal/mol. b) Example of one of the unstable conformer states probed that exploded, despite the bias potential used during the optimisation.

**Figure S13:**
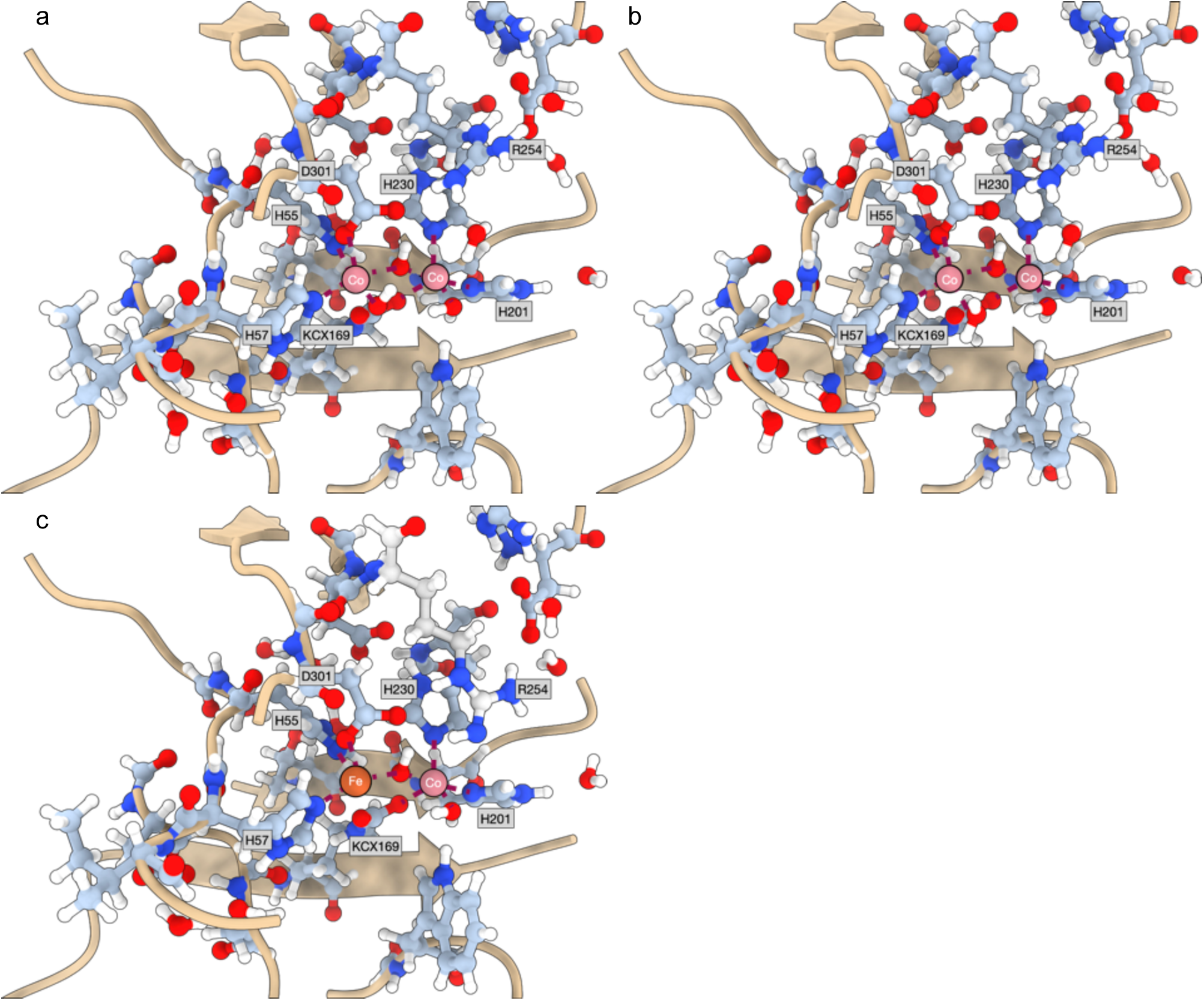
The three states of 3wml. Complete characterisation of the states in 3wml. a) The Co-Co state with a hydroxyl group coordinated to the leftmost cobalt ion. b) The Co-Co state with a coordinated water molecule. c) The Fe-Co state, with a change in conformation of the residue R254. This residue is furthermore marked with a different colour.

**Figure S14:**
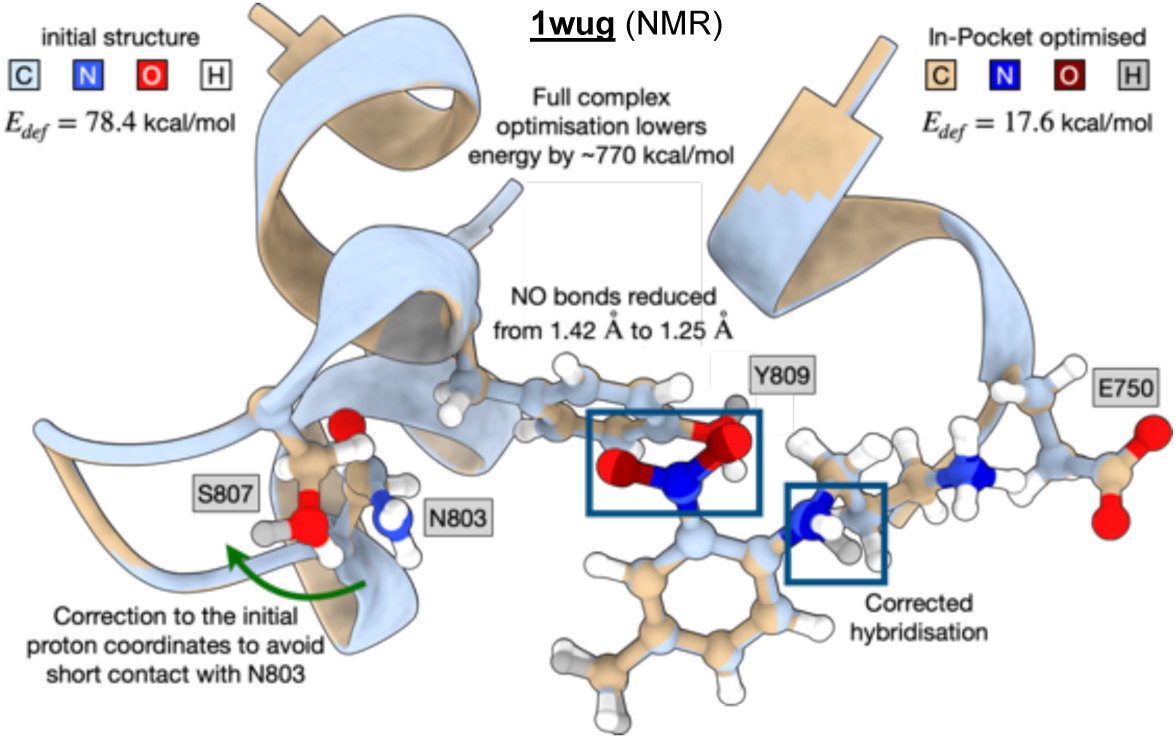
IPA refinement of the NMR structure 1wug. The initial ligand’s deformation energy of 78.4 kcal/mol can be reduced to 17.6 kcal/mol using IPA, resulting from corrected N-O bond distances in a nitro-group and fixes in atomic hybridisations. Structural distortions in nitro groups are pervasive in PDB-deposited models. In addition to bond length discrepancies, asymmetric N-O bond distances are frequently observed within one monomer, which are inconsistent with the electronic structure of nitro groups, even in the anisotropic environment of a protein’s pocket. To further investigate these distortions, we analysed the 1wug protein-ligand complex (CCD-ID: NP1),^44^ which features a nitro-containing ligand bound to a bromodomain (**Figure S14**). The IPA refinement significantly reduced N-O bond distortions while optimising the hydrogen-bonding network. These structural corrections translated into substantial energetic improvements: the overall system stabilised by 770.0 kcal/mol due to a more favourable network of interactions, while the ligand-specific deformation energy dropped by 60.8 kcal/mol. Notably, IPA also corrected the hybridisation states of the aniline-like nitrogen, a required adjustment since classical protonation algorithms rely exclusively on geometric heuristics. These refinements emphasise the importance of QM corrections in ensuring chemically accurate ligand representations within protein-ligand complexes.

**Figure S15:**
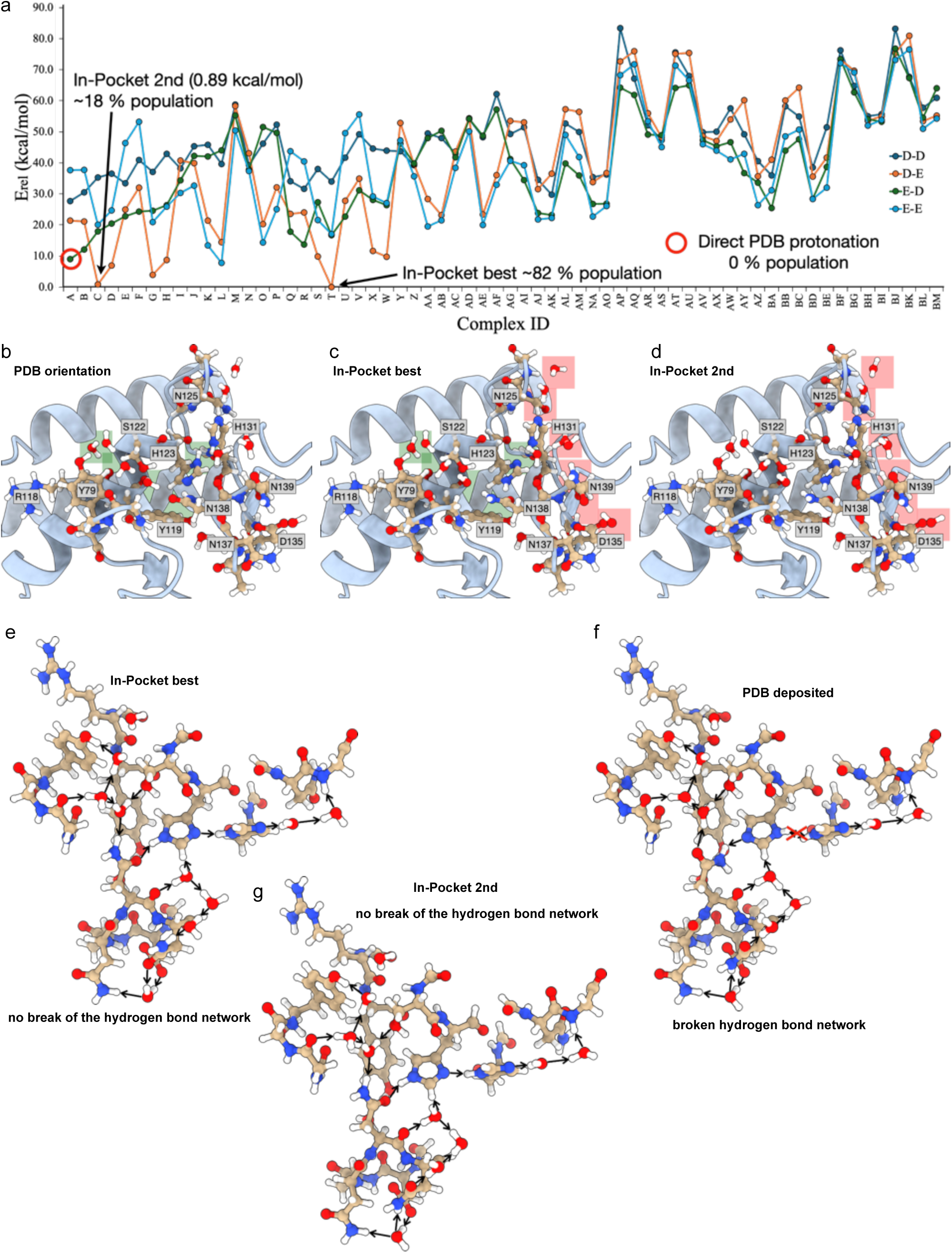
Alternative hydrogen-bond networks in the PDB 1cem. a) The relative energy of each network state identified for the selected fragment of the PDB structure 1cem. For each histidine protonation pattern (δδ = D-D; δε = D-E; εδ = E-D; εε = EE), 64 orientational conformers were identified and refined with IPA. Depending on the protonation pattern selected for the histidine residues, the PDB-deposited structure is at least 10.0 kcal/mol higher in energy than the lowest energy state, indicating that this structure is thermodynamically irrelevant to describe the system. A direct PDB protonation corresponds to proton addition according to the PDB-deposited topology, following a δ-protonation of histidines. IPA’s fragment mode improves b) the initial PDB-deposited conformer. The green shaded areas locate the major differences recorded between the hydrogen-bond refined PDB structure and the c) IPA minimum energy structure. d) The second-lowest energy system identified by IPA. The red-shaded areas show differences with respect to the IPA energy minimum. The second most stable structure is about 0.89 kcal/mol less favourable than the minimum found. e) The hydrogen-bond network of the lowest energy chemical state identified for the 1cem fragment we analysed. An uninterrupted network of 18 hydrogen bonds justifies the stability of this conformer. f) The hydrogen-bond network for the PDB-deposited chemical state shows an interruption due to a proton clash between the histidine residues. This structure is thermodynamically irrelevant to describe the system. g) The second most stable IPA identified state also shows an uninterrupted network of 18 hydrogen bonds, though slightly less favourable than the energy minimum. To assess IPA’s ability to analyse extensive hydrogen-bond networks in biological systems, we examined 256 proton-network orientations in the PDB model 1cem^45^ (**Figure S15**): residues H123 and H131 may be δδ, δε, εδ, or εε protonated, yielding 64 possible structures obtained by combinatorially flipping His, Asn, and Gln residues. Inherently, the surrounding water network adapts to side chain relative orientations. Of all conformations generated, only structures A–δδ, δε, εδ, and εε–are consistent with the deposited PDB structure. Our calculations reveal that the PDB structure with δδ His protonation is nearly 30.0 kcal/mol higher in energy than the lowest-energy conformer (**Figure S15a**). Boltzmann statistics indicate that a naïve, direct protonation of the structure according to the PDB topology is thermodynamically irrelevant. The elevated energy results from a proton clash that disrupts the hydrogen-bond network (**Figures S15b** and **S13b**). An εδ His protonation moderately improves the system’s energetics, yet it remains 10.0 kcal/mol above the lowest-energy state and is similarly biologically irrelevant. The most stable conformation was identified after three Asn flips (N125, N138, and N139) and adopting a δε His protonation (T conformer, **Figures S15a,c**, and **S15e**). This state accounts for approximately 82% of the conformer population. The second most populated conformer, featuring a single N139 flip from the PDB structure with δε His protonation (**Figures S15a,d**, and **S15g**), lies 0.9kcal/mol above the minimum energy state and represents ∼18% of the total population. To benchmark further, we used MolProbity^46^ and WHAT_CHECK^47^and observed agreement with the IPA results.

**Figure S16:**
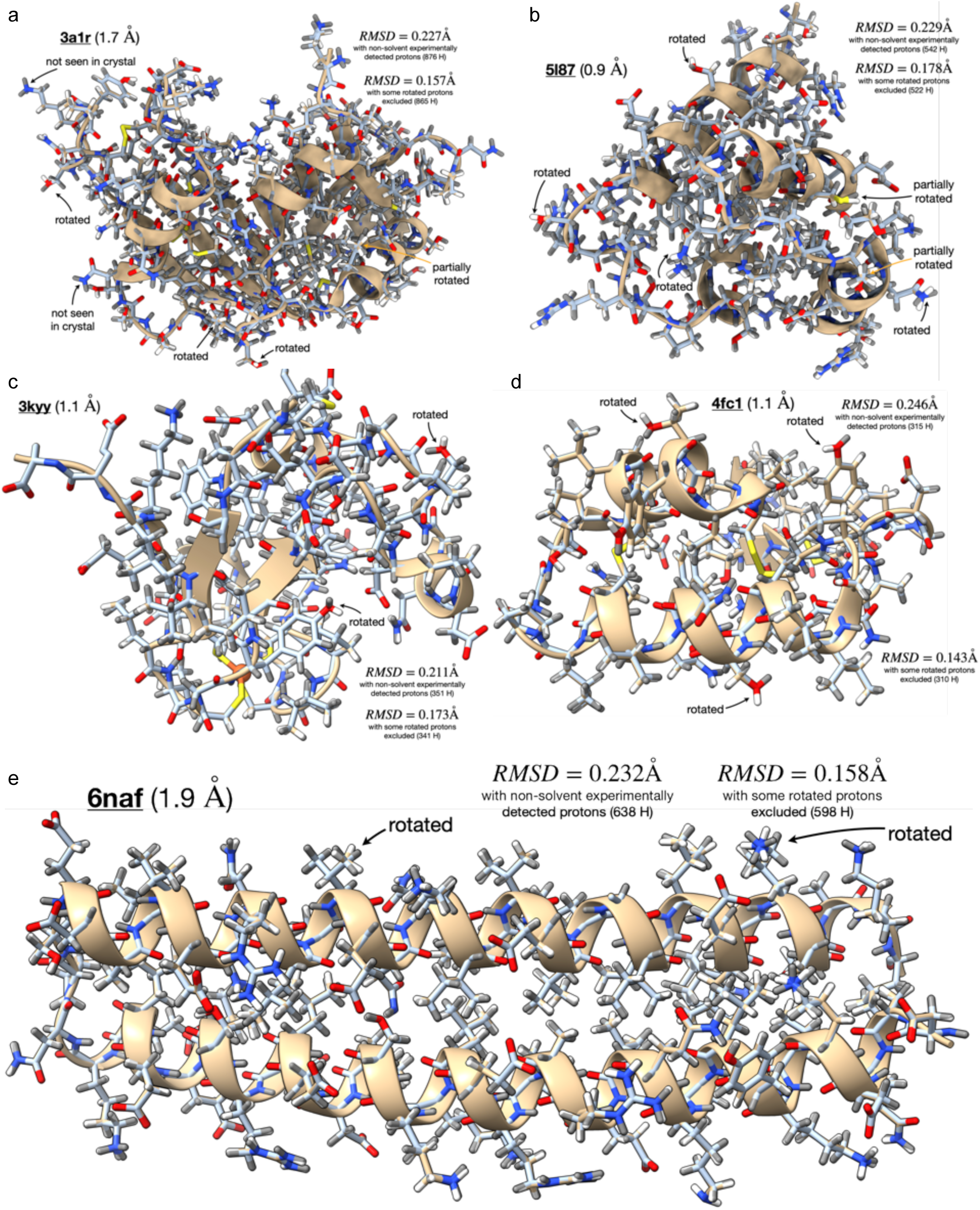
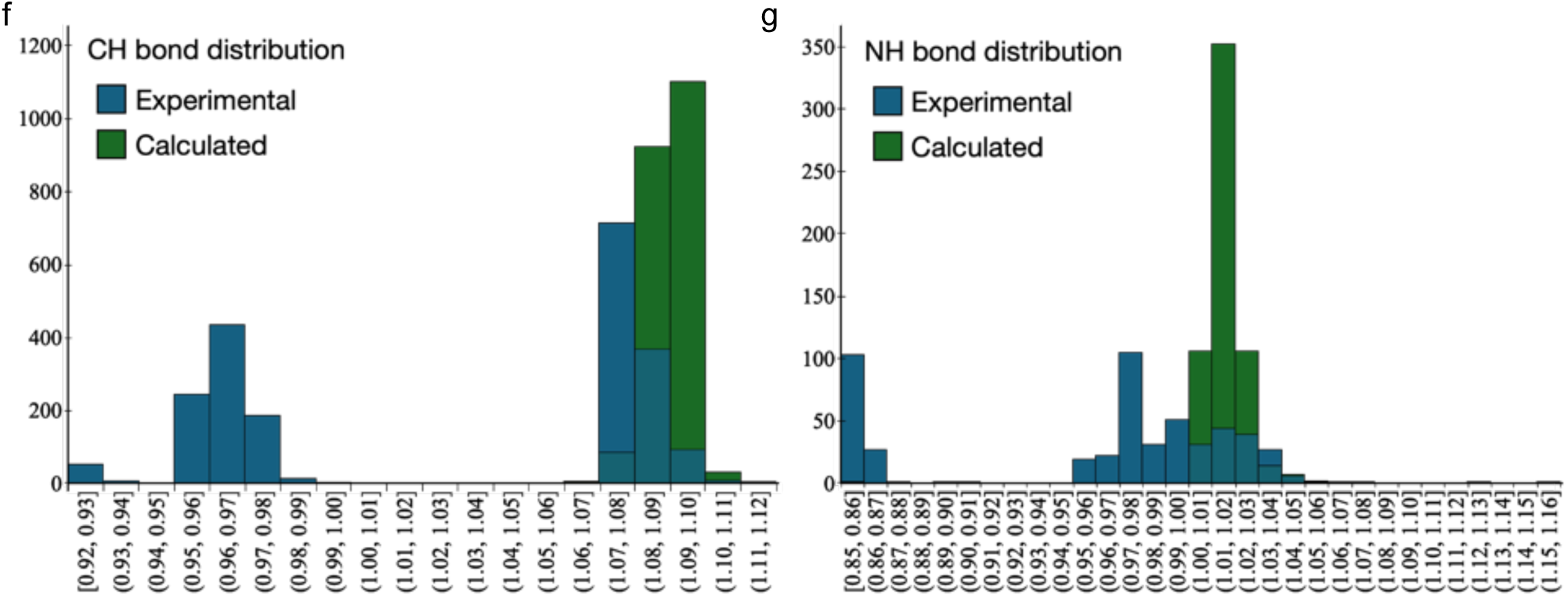
Additional IPA validation through proton-refined coordinates. Validation using the PDBs a) 3a1r,^48^ b) 5l87,^49^ c) 3kyy,^50^ d) 4fc1,^51^ and e) 6naf.^52^ These cases compare the IPA-optimised proton network against experimentally declared proton coordinates. Besides a few rotamers observed on the protein’s solvent-exposed surface, the two sets of proton coordinates are in great agreement. The structures, spanning various resolutions, were re-protonated to introduce randomness in initial proton placements, while all non-hydrogen atoms were preserved to best reflect the typically experimentally observed system. The accuracy of IPA-optimised structures was assessed by computing the root-mean-square deviation (RMSD) between initial and final proton coordinates, yielding values of 0.21–0.25Å. This analysis included only protein and, where present, small-molecule proton coordinates, excluding solvent and crystallisation additives. A closer inspection reveals that major deviations arise primarily from freely rotating groups, such as terminal methyl or ammonium moieties. Additionally, some solvent-exposed hydroxyl groups (*e.g.*, tyrosine) remained flipped by 180° due to their randomised starting orientations. Correcting these orientations to match experimental data would require overcoming an activation barrier, which is unfeasible for geometry optimisation algorithms. Excluding these protons from the RMSD calculation reduced the experimental-calculated RMSDs to 0.14-0.18Å, highlighting the precision achievable with IPA. f) Comparing the IPA-optimised CH bond distance distributions against the experimentally observed ones. The former shows a unimodal distribution, whereas the latter is multimodal. g) Comparing the IPA-optimised NH bond distance distributions against the experimentally observed ones. Again, the former shows a unimodal distribution, whereas the latter is multimodal.

**Figure S17:**
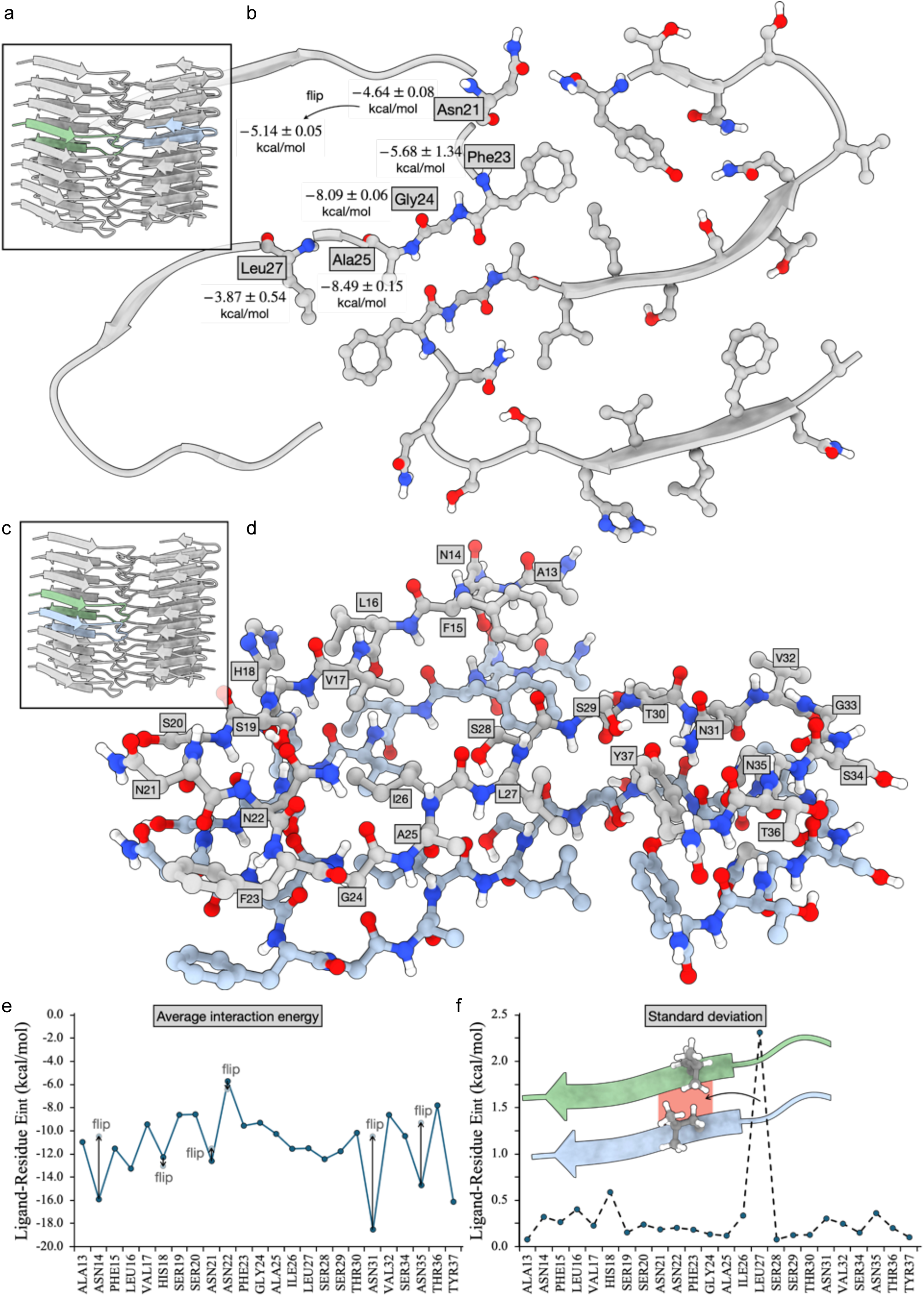
IPA validation of the protein-protein interaction interfaces of the amyloid fibril cryo-EM structure 6y1a.^53^. The experimental structure comprises two columns of stacked peptides, generating two types of protein-protein interfaces (PPIs): a-b) A small interface formed by equally ranked peptides from different columns; c-f) A large interface generated by adjacent peptides within the same column. a) Overall architecture of the stacked assembly, with the green and blue ribbons highlighting one of the small interaction interfaces examined. IPA analysis was extended to all peptide pairs forming a small interface. b) Detailed structural model of one small interface, including corresponding energetics and standard deviations. The interface is stabilised by five principal contacts involving Asn21, Phe23, Gly24, Ala25, and Leu27. Note, however, that not all of these interactions fall within the 4 Å threshold radius used for the calculations; for example, contacts with Gly24 and Leu27 are conserved in four of the eight models analysed. Overall, most contacts show limited standard deviation (σ; ± values in block b), except those involving Phe23 (σ = 1.34 kcal mol^−1^) and Leu27 (σ = 0.54 kcal mol^−1^). Variability in Phe23 arises because, in some models, Phe23 approaches Tyr37 too closely, resulting in a mild steric clash between protons. Deviations in Leu27 interactions reflect its coupling with Phe23. The energetic analysis indicates that flipping Asn21 slightly improves the interaction network. c) Placement of the large interaction interface within the context of the fibril stacked structure. d) Detailed structural model of the large interface with all residues named using one-letter amino acid codes. The main statistics for all ligand-residue interaction energies observed in the large interface, represented as e) average values and f) their respective standard deviations (σ). All average interaction energies are strong, reflecting the high incidence of hydrogen bonds. Analysis of these energies indicates that flipping key Asn and His residues generally weakens the interaction interface, except for His18 and Asn22, where the improvement upon flipping is minor. In contrast to the short-interface analysis, flipping Asn21 results in a slight energetic deterioration. Examination of the standard deviations reveals uniformly low variability across most residues, except for Leu27, which exhibits a higher deviation due to steric clashes arising from Leu27–Leu27 stacking between consecutive peptides.

**Figure S18:**
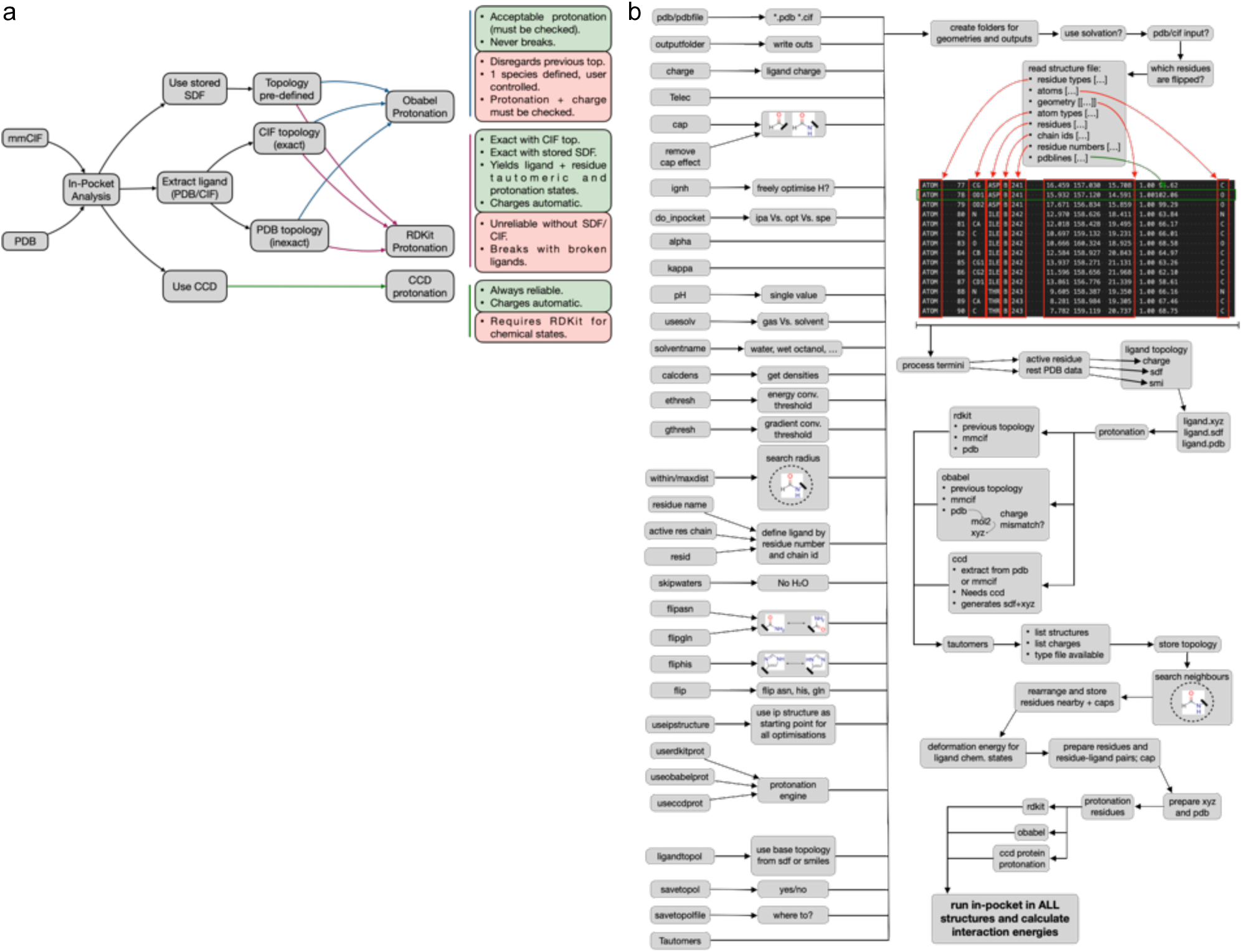
Schematic Representations of the IPA algorithms and an overview of IPA options. a) Schematic representation of how the IPA algorithm works. It is possible to start the process using a PDB or a mmCIF file. Different algorithms may be used for protonating structures and to proceed with further processing. Each algorithm’s advantages (green) and disadvantages (red) are indicated next to it. b) Schematic overview of how IPA works and an overview of all options available.

